# Quantification of morphological, functional, and biochemical features of H9c2 rat cardiomyoblast retinoic acid differentiation

**DOI:** 10.1101/2024.12.10.627787

**Authors:** Nicole S. York, Joel E. Rivera, Mohammadreza Rahmani Manesh, K’sana Wood Lynes-Ford, Rory Smith, Leigh E. Wicki-Stordeur, Laura T. Arbour, Leigh Anne Swayne

**Affiliations:** Division of Medical Sciences, University of Victoria, Victoria, British Columbia, Canada; Department of Medical Genetics, University of British Columbia, Victoria, British Columbia, Canada; Island Medical Program, University of British Columbia, Victoria, British Columbia, Canada

**Keywords:** H9c2, differentiation, retinoic acid, morphology, spontaneous calcium transients

## Abstract

Cell culture models enable advancement in our understanding of heart development and heart disease. The H9c2 rat ventricular cardiomyoblast cell line can be differentiated, recapitulating certain components of cardiomyocyte development; however, morphological, functional, and biochemical changes are rarely investigated in parallel, thereby limiting fulsome understanding. We therefore characterized morphology, Ca^2+^ handling, and gene expression after five days (5 *days-in-vitro*, DIV5), and fourteen days (DIV14) of exposure to differentiation stimuli, consisting of retinoic acid and low serum. We observed several morphological changes with increasing time in differentiation, including increased mean length, area, eccentricity, and multinucleation, as well as an increase in actin clusters. Spontaneous Ca^2+^ transients in differentiated H9c2 cells had decreased frequency and synchronicity compared to those observed in primary cardiomyocytes. Additionally, key cardiomyocyte cytoskeletal proteins and ion channel transcript and protein expression levels changed significantly with differentiation, in alignment with changes normally observed in cardiomyocyte development. Our findings suggest H9c2 cells may be used as a high throughput screening model to investigate cardiomyocyte biology in certain developmental contexts with relevance to heart health and disease.

**Highlights:**

- We quantified morphology, Ca^2+^ handling, and gene expression changes in rat ventricular cardiomyoblast H9c2 cells across retinoic acid differentiation.
- We compared multiple differentiation time points (mainly DIV0, DIV5, and DIV14) and observed time-dependent changes in all 3 aspects. We observed several morphological changes with increasing time in differentiation, including increased mean length, area, eccentricity, and multinucleation, as well as an increase in actin clusters. We observed the onset of spontaneous Ca^2+^ transients with differentiation, with further changes in subpopulations with increasing time in differentiation. Finally, we observed multiple changes in gene expression consistent with cardiomyocyte differentiation.
- We summarize these parallel time-point dependent changes in morphology, Ca^2+^ handing, and gene expression to enable other researchers to optimize their study design.

## Introduction

Cardiomyocyte-like cell culture models, including immortalized cell lines such as the H9c2 rat ventricular cardiomyoblast cell line, human induced pluripotent stem cell (iPSC)-derived cardiomyocytes, and rodent primary cardiomyocyte cultures play key, complementary roles in modeling cardiomyocyte development. Together, their coordinated use could enable fulsome modelling of the cellular changes in health and disease contexts. Each of these models has key strengths and weaknesses facilitating the study of different aspects of cardiomyocyte development and viability. The H9c2 cardiomyoblast cell line can be differentiated with low serum and retinoic acid to express phenotypes associated with cardiomyocytes [1–3]. As an immortalized cell line, H9c2 cells convenient to work with, suggest retinoic acid-differentiated H9c2 cells could be an attractive model to study aspects of cardiomyoblast and/or cardiomyocyte biology (Figure 1). However, there remain significant gaps in the quantitative changes in these cells across different cellular parameters altered by differentiation, that hinder their use. To this we measured changes in H9c2 cell morphology, Ca2+ handling, and gene expression across retinoic acid differentiation, to better enable their use as a model for cardiomyoblast and/or cardiomyocyte biology in health and disease contexts.

**Figure 1.**
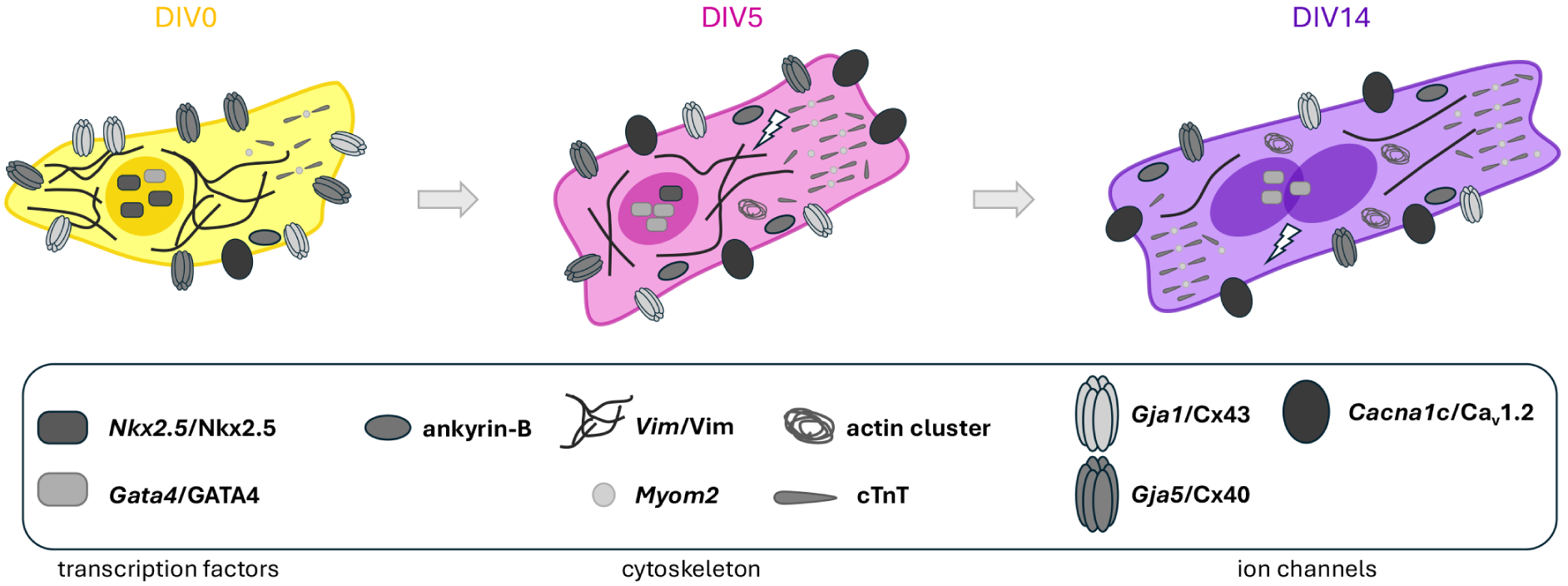
H9c2 cells display key markers of cardiomyocytes. H9c2 cells undergo a variety of genetic changes during H9c2 cell differentiation, upregulating key cardiomyocyte genes and proteins such as Cacna1c (Cav1.2), Tnnt2 (cTnT), and Myom2 (Myomesin-2).

The H9c2 cell line was first isolated from the lower half of a rat ventricle at embryonic day 13 [4]. Following selective passage to enrich for cardiomyoblasts, the cells ceased beating but maintained other key cardiomyoblast characteristics [4], such as their morphology, ultrastructure, and presence of specific ion channels [1,4–6]. Early work showed that retinoic acid differentiation increased Ca_v_1.2 (α_1C_)-based L-type Ca^2+^ channel expression [1], which are enriched in and important for generating the electrical and mechanical properties of cardiomyocytes [7], and cardiac troponins [2]. Several additional studies have investigated the morphological and biochemical aspects of differentiated H9c2 cells [2,3], but functional studies are rare [1,8]. H9c2 cell differentiation is associated with elongation of cell shape [3,6–8], but this has not been fully quantified [1–4].

Consequently, the amplitude and extent of changes in basic morphological properties associated with differentiation is unclear. To date, characterization of H9c2 cell intracellular Ca^2+^ handling has been limited to stimulation of undifferentiated cells [8–10], but the impact of differentiation on baseline H9c2 cell Ca^2+^ handling properties is unknown. H9c2 cell differentiation is associated with increased expression of genes encoding cardiomyocyte markers, such as cardiac troponins, and genes involved in Ca^2+^ handling and mitochondrial metabolism [2]. Although H9c2 cell differentiation-associated gene expression changes are better defined [2,3,8], they have rarely been compared with morphological and/or functional aspects in parallel. Furthermore, early changes (e.g., 5 hours to 3 days), which could be driving various signaling processes, have not been thoroughly investigated. The amplitude and timing of gene expression changes relative to morphological and Ca2+ handling changes is rarely studied in parallel, making it difficult to identify possible connections.

To this end, we quantified morphology, Ca^2+^ handling, and gene expression changes across H9c2 cell retinoic acid differentiation in parallel, comparing mainly three time points, DIV0, DIV5, and DIV14. We used confocal and super-resolution microscopy, intracellular Ca^2+^ imaging, as well as RT-qPCR and Western blotting to assess these changes. We observed several morphological changes with increasing time in differentiation, including increased mean length, area, eccentricity, and multinucleation, as well as an increase in actin clusters. We observed the onset of spontaneous Ca^2+^ transients with differentiation, with further changes in subpopulations with increasing time in differentiation. We also observed multiple changes in gene expression consistent with cardiomyocyte differentiation, such as increased *Myom2*, *Cacna1c*, GATA4, and cTnT expression. Finally, we compiled and summarize these changes according to DIV to enable optimal use of differentiated H9c2 cells in modelling aspects of cardiomyoblast and/or cardiomyocytes in health and disease contexts.

## Results

### Differentiated H9c2 cells become larger, multi-nucleated, and develop localized actin intensities

During cardiomyocyte differentiation, cells undergo changes in their overall shape that are key for their function [11–13]. Prior work commonly differentiated H9c2 cells up to DIV5 [2,3] yet more recent work has extended differentiation to DIV14 [8]. We compared an array of gross morphology features of undifferentiated (DIV0), DIV5- and DIV14- differentiated H9c2 cells by analyzing confocal images of F-actin (phalloidin) and nuclei (Hoechst) (Figure 2A) with a custom MATLAB script (available on github), refer to Table 1 for statistical analyses. Average cell length was significantly greater at DIV14 compared to DIV0, with corresponding right shifts in the frequency distribution plot of cell length for both DIV5 and DIV14 cell populations (Figure 2Bi, Ci). Cell area also significantly increased at DIV14 compared to both DIV0 and DIV5 cells, and the frequency distribution plot of cell area exhibited right shifts for both DIV5 and DIV14 cell populations (Figure 2Bii, Cii). Cell eccentricity increased at DIV5 compared to DIV0, with no further change evident at DIV14 (Figure 2Biii). Corresponding right shifts were evident in the frequency distribution plot of cell eccentricity for both DIV5 and DIV14 cell populations (Figure 2Ciii). The proportion of multinucleated cells was also drastically increased at DIV14 compared to DIV0 (Figure 2Biv, Civ). We found no significant changes in mean cell perimeter across differentiation, however the frequency distribution plot for cell perimeter did show some right shifting of DIV5 and DIV14 cell populations (Figure S1). Together these data indicate that differentiated H9c2 cells are, on average, longer, larger, more eccentric (less circular) and more likely to be multinucleated than undifferentiated DIV0 cells. Moreover, these changes are more apparent at the DIV14 differentiation timepoint than at DIV5, suggesting that extending the differentiation time course further enhances cardiomyocyte-like phenotypic changes.

**Figure 2.**
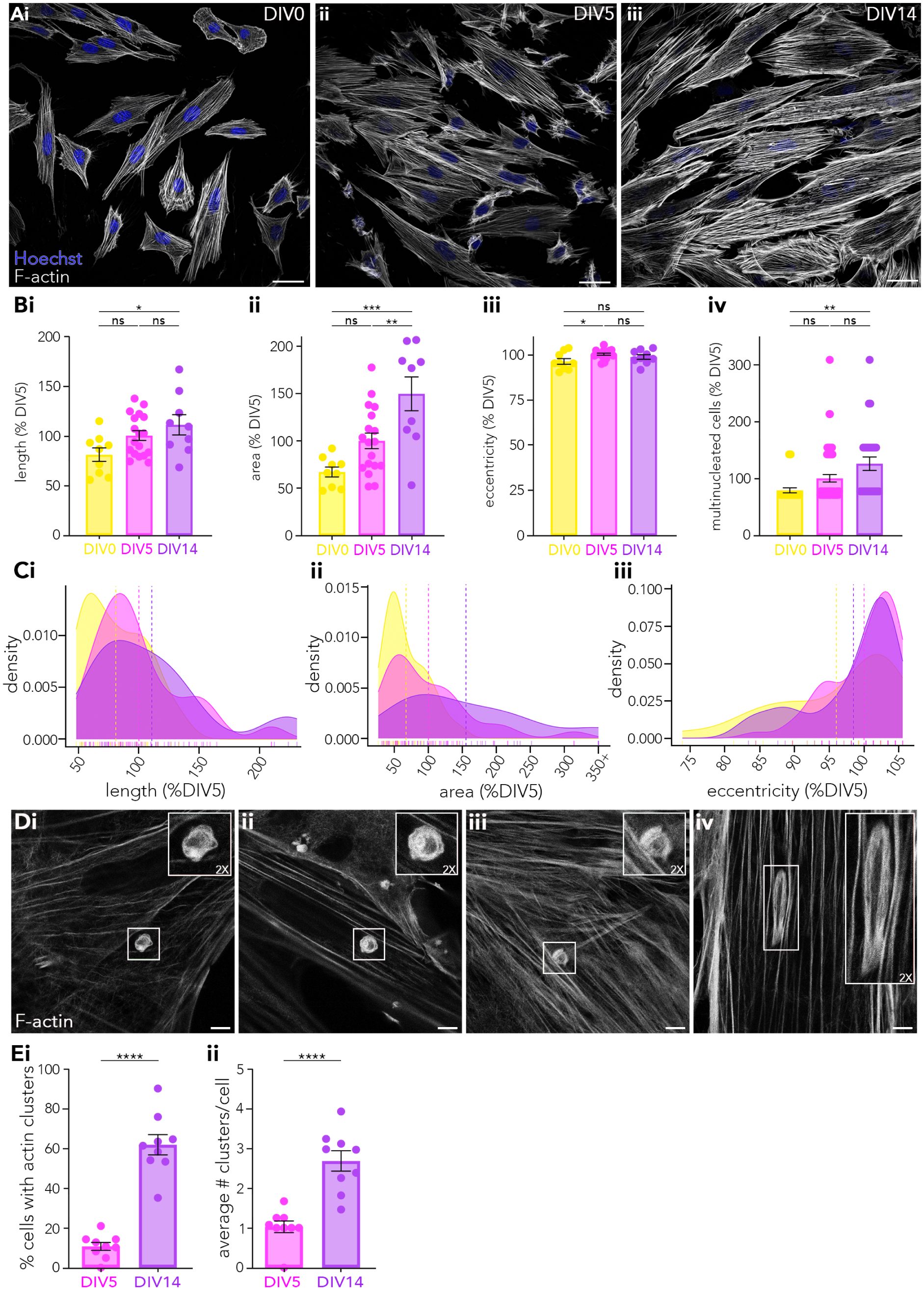
Differentiated H9c2 cells become larger, multi-nucleated, and develop localized actin intensities at DIV14. H9c2 cell morphology at (Ai) DIV0, (Aii) DIV5, and (Aiii) DIV14 were analyzed using the F-actin stain Phalloidin (white) and nuclei were counted via Hoechst staining (blue). Length, perimeter, area, eccentricity, and the number of nuclei per cell were measured using a custom MATLAB script which resulted in significant increases in (Bi) length, (Bii) area, (Biii) eccentricity, and (Biii) percentage of multinucleated cells. Density distribution plots show cell (Ci) length, (Cii) area, and (Ciii) eccentricity at DIV5 display larger values than at DIV0. (Di-iii) Circular and (Div) elliptical actin-positive clusters were present in many of the cells. Insets show STED resolved clusters at 2X magnification. (Ei) Quantification of the percent of cells displaying actin clusters is higher at DIV14 than DIV5. (Eii) The number of actin clusters per cell also is increased at DIV14. (B, E) Data is presented as mean ± SEM and (C) display probability density plots, indicating how likely a particular value is to occur. Results are graphed as Yellow = DIV0, Magenta = DIV5, Purple = DIV14. (A) Scale bar = 40 µm. (D) Scale bar = 5 µm, inlays were zoomed in by a factor of 2. N = 3 replicates, n = 27 cells for all groups (3 cells from each of the 3 FOV were selected). ****, p < 0.0001; ***, p < 0.001; **, p < 0.01; *, p < 0.05; ns, p > 0.05 by (B) one-way ANOVA and post-hoc Tukey’s multiple comparison and normalized to DIV5, or by (E) Independent T-test, see Table 1 for detailed statistics

**Table 1.** Summary statistics of H9c2 cell morphology analysis results in significant increases to length, area, eccentricity, and multinucleation across differentiation. Minimum (Min) and maximum (Max) values for each condition as well as the mean ± SEM are listed. A one-way ANOVA was performed, if this provided a significant result, then the Tukey multiple comparisons test was used to determine which groups were significant. N=9 FOV for DIV0, DIV14, and N=18 FOV for DIV5 each from a total of 3 passages. All values are normalized to DIV5.

| parameter | timepoint | min (%) | max (%) | mean ± SEM | p-value ANOVA | p-value Tukey | | |
| --- | --- | --- | --- | --- | --- | --- | --- | --- |
|  |  |  |  |  |  | 0-5 | 0-14 | 5-14 |
| normalized length | DIV0 | 55.5 | 114.4 | 80.8% ± 6.7 | | | | |
| | DIV5 | 72.9 | 137.1 | 100% ± 4.9 | 0.0321<br>*** | 0.1258<br>"ns" | 0.0278<br>*** | 0.5083<br>"ns" |
| | DIV14 | 68.1 | 166.3 | 110.7% ± 10.2 | | | | |
| normalized perimeter | DIV0 | 36.2 | 137.1 | 77.2% ± 10.5 | | | | |
| | DIV5 | 47.4 | 177.0 | 100% ± 8.5 | 0.2392<br>"ns" | 0.3290<br>"ns" | 0.2567<br>"ns" | 0.9122<br>ns" |
| | DIV14 | 47.4 | 186.0 | 106.4% ± 16.3 | | | | |
| normalized area | DIV0 | 47.2 | 92.0 | 67.1% ± 5.3 | | | | |
| | DIV5 | 51.7 | 177.5 | 100% ± 8.3 | 0.0002<br>**** | 0.0934<br>"ns" | 0.0001<br>**** | 0.0072<br>*** |
| | DIV14 | 53.2 | 206.5 | 149.7% ± 17.9 | | | | |
| normalized eccentricity | DIV0 | 90.1 | 103.4 | 96.0% ± 1.6 | | | | |
| | DIV5 | 94.6 | 105.2 | 100% ± 0.6 | 0.0302<br>*** | 0.0231<br>*** | 0.3077<br>"ns" | 0.5412<br>"ns" |
| | DIV14 | 91.5 | 103.4 | 98.5% ± 1.3 | | | | |
| normalized multinucleation | DIV0 | 71.1 | 142.1 | 79.0% ± 4.4 | | | | |
| | DIV5 | 71.1 | 308.6 | 100% ± 6.5 | 0.0018<br>*** | 0.1446<br>"ns" | 0.0012<br>*** | 0.0578<br>"ns" |
| | DIV14 | 77.1 | 308.6 | 125.7% ± 11.8 | | | | |

Notably, we observed that DIV14 differentiated H9c2 cells commonly exhibited localized actin intensities, or “clusters” of F-actin (phalloidin+) (Figure 2Aiii) like those previously reported in other cell types [14–16]. We used stimulated emission depletion (STED) super-resolution microscopy to further resolve these F-actin clusters, which were circular to ellipsoid in nature (Figure 2D). At DIV14, 61.8% ± 5.1 of cells showed actin clusters compared to 10.8% ± 2.0 of cells at DIV5 (unpaired t-test, p<0.0001; Figure 2Ei). Of the cells that exhibited actin clusters, the average number of clusters per cell increased between these time points (DIV5, 1.0 ± 0.1 clusters/cell *vs*. DIV14, 2.7 ± 0.3 clusters/cell; unpaired t-test, p<0.0001; Figure 2Eii).

Together these results suggest that retinoic acid differentiation increases cell length, cell area, eccentricity, and the proportion of multinucleated cells, and induces the formation of circular/elliptical F-actin clusters.

### Differentiated H9c2 cells develop spontaneous Ca^2+^ transients similar in amplitude to primary cardiomyocytes

Intracellular Ca^2+^ handling changes dynamically across the development of cardiomyoblasts into cardiomyocytes [17,18]. But Ca^2+^ handling during H9c2 cell differentiation has not been thoroughly characterized, making it difficult to begin to relate changes in morphology and gene expression to functional outcomes. To this end, with live confocal imaging of Fluo4-AM [19] and previously published MATLAB scripts [20], we examined spontaneous Ca^2+^ transients in undifferentiated and differentiated H9c2 cells and compared DIV5 and DIV14 differentiated H9c2 cells with neonatal primary mouse cardiomyocytes (Figure 3A). At DIV0 we did not observe readily detectable spontaneous Ca^2+^ transients (not shown) and therefore, DIV0 was not included in subsequent analyses. To determine the extent to which differentiation brings H9c2 cells towards cardiomyocytes’ Ca^2+^ handling, all metrics were compared to those of primary cardiomyocytes. Mean transient (spike) amplitudes (change in fluorescence from baseline to peak) of all H9c2 cell differentiation conditions were similar to primary cardiomyocytes (Figure 3Bi, Table 2). However, mean spike frequency (spikes per minute) and synchronicity (number of spikes occurring at the same time that exceed a threshold based off the mean intensity profile in all the cells in the field of view (FOV)) were significantly lower in all differentiated H9c2 cell conditions compared to primary cardiomyocytes (Figure 3Bii,iii, Table 2). Distribution plots of frequency and synchronicity data displayed right shifts of the DIV14 population compared to DIV5, suggesting a further shift of DIV14 cells closer to the primary cardiomyocyte phenotype.

**Figure 3.**
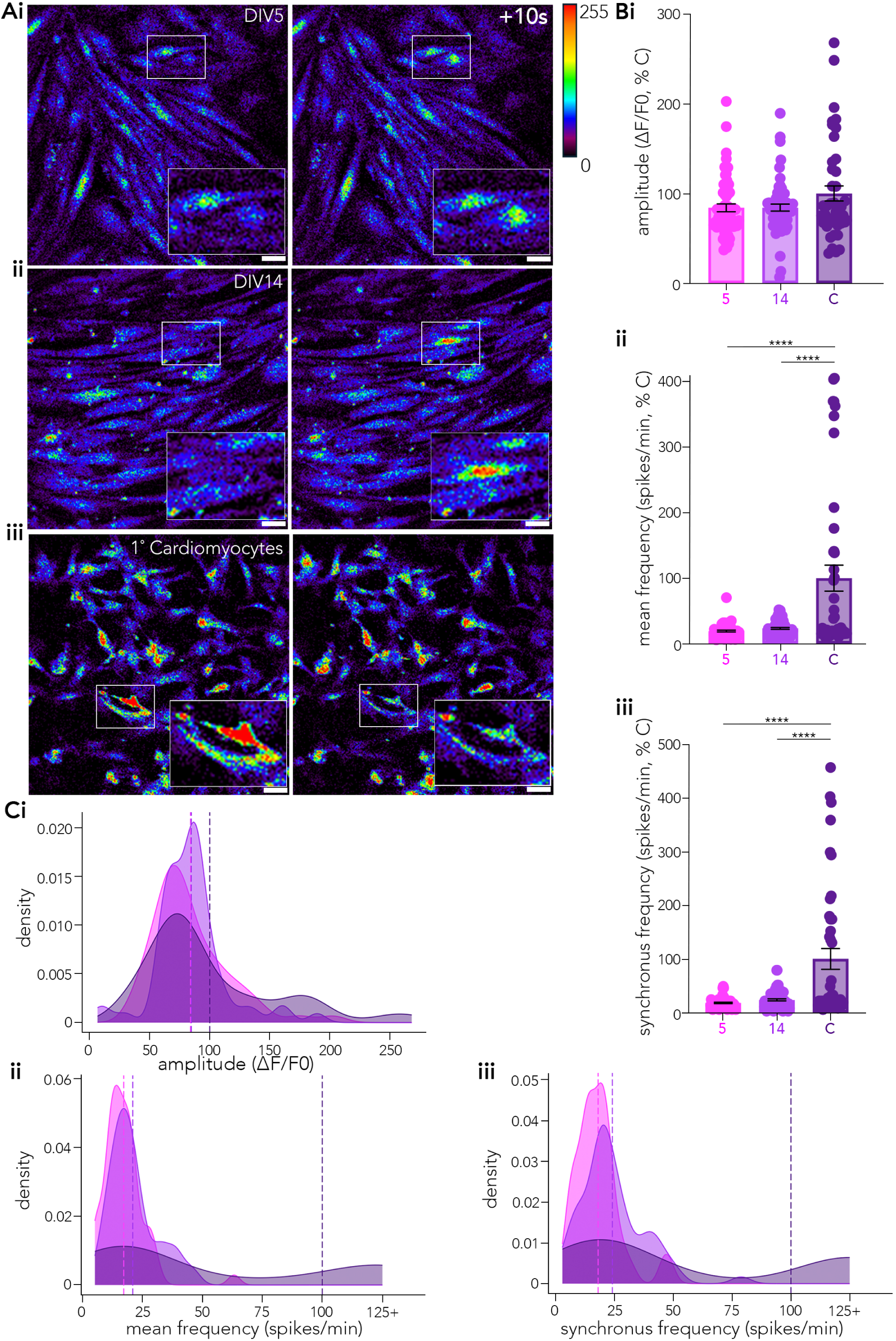
Differentiated H9c2 cells exhibit spontaneous Ca^2+^ transients, subpopulations of which exhibit characteristics nearing those of primary cardiomyocytes. Ca^2+^ imaging of differentiated H9c2 cells was performed using fluo-4 AM. Stills 10 seconds apart from 2 min (Ai) H9c2 DIV5, (Ai) H9c2 DIV14, or 1 min (Aii) primary cardiomyocytes time series show spontaneous changes in [Ca^2+^]i. See supplemental Videos 1-3 for corresponding live imaging recordings. Inlays highlight active cells in the field of view (FOV) undergoing a spike in [Ca^2+^]i. H9c2 cells have similar (Bi) spike amplitude to primary cardiomyocytes while (Bii) frequency and (Biii) synchronous firing rate are significantly different. Density distribution plots show the probability of a value in each group for (Ci) spike amplitude, (Cii) frequency, and (Ciii) synchronous firing rate highlighting right shifts at DIV14 for (Cii) frequency and (Ciii) synchronous firing rate. DIV5 (N=56 FOV from 7 passages), DIV14 (N =60 FOV from 6 passages), primary cardiomyocytes (C; N = 43 FOV from 5 cultures). Data is presented as mean ± SEM. Results are graphed as Magenta = DIV5, Purple = DIV14, Violet = C. Scale bar = 40 µm. ****, p < 0.0001; unlabeled, p > 0.05 by one-way ANOVA and post-hoc Tukey’s multiple comparison. Data is normalized to primary cardiomyocytes, see Table 2 for detailed statistics.

**Table 2.** Summary statistics of H9c2 cell spontaneous calcium dynamics. Minimum (Min) and maximum (Max) values for each condition as well as the mean ± SEM are listed for each parameter. DIV5 (N=56 FOV from 7 passages), DIV14 (N=60 FOV from 6 passages), primary cardiomyocytes (N=43 FOV from 5 cultures). A one-way ANOVA was performed, if this provided a significant result, then the Tukey multiple comparisons test was used to determine which groups were significant. All values are normalized to primary cardiomyocytes (C).

| parameter | timepoint | min (%) | ax (%) | mean ± SEM | p-value ANOVA | p-value Tukey | | |
| --- | --- | --- | --- | --- | --- | --- | --- | --- |
|  |  |  |  |  |  | 5-14 | 5-C | 14-C |
| normalized amplitude | DIV5 | 37.1 | 202.3 | 84.1% ± 4.3 | | | | |
| | DIV14 | 6.8 | 189.1 | 84.2% ± 3.9 | 0.0789<br>"ns" | 0.9999<br>"ns" | 0.1138<br>"ns" | 0.1099<br>"ns" |
| | C | 33.4 | 267.6 | 100% ± 8.5 | | | | |
| normalized frequency | DIV5 | 6.0 | 70.4 | 19.3% ± 1.3 | | | | |
| | DIV15 | 7.7 | 51.9 | 23.5% ± 1.4 | <0.0001<br>"*****" | 0.9413<br>"ns" | <<br>0.0001<br>"*****" | <<br>0.0001<br>"*****" |
| | C | 8.9 | 404.8 | 100% ± 19.8 | | | | |
| normalized synchronous frequency | DIV5 | 5.4 | 48.9 | 18.0% ± 1.3 | | | | |
| | DIV14 | 2.7 | 78.8 | 23.7% ± 1.7 | <0.0001<br>"*****" | 0.8639<br>"ns" | <<br>0.0001<br>"*****" | <<br>0.0001<br>"*****" |
| | C | 5.4 | 456.5 | 100% ± 19.4 | | | | |

Together these results suggest that spontaneous Ca^2+^ transients develop with differentiation of H9c2 cells and are comparable in amplitude to those of primary cardiomyocytes. Notably, while the longer differentiation timepoint (DIV14) does not overtly alter H9c2 cell Ca^2+^ transients compared to the DIV5 timepoint, there may be subpopulations of DIV14 cells that are shifted closer towards a primary cardiomyocyte-like phenotype.

### Differentiated H9c2 cells exhibit gene expression changes consistent with cardiomyocyte differentiation

In parallel to our assessment of changes in H9c2 cell morphology and Ca^2+^ handling associated with differentiation, we probed transcript (RT-qPCR) and protein (Western blotting) expression levels of genes involved in transcription, cytoskeleton, ion flux regulation, and signal transduction (Figure 4A, see Table S1 for brief descriptions of functions and references). Many of these gene expression changes have been previously described in H9c2 cell differentiation (Rbp2, Tnnt2, Myom2, Pde4a [2]). We also looked at additional gene expression changes based on the cardiomyocyte literature (Nkx2.5, a-actinin). For select transcript changes we confirmed protein expression with Western blotting (cTnT, GATA4, Nkx2.5, β-actin, Cx43, vimentin), and in some cases used only used Western blotting (a-actinin, vinculin, ankyrin-B). We used two types of comparisons. In the first approach, to identify increases or decreases in gene expression associated with H9c2 cell differentiation, we quantified the levels of select transcripts and proteins at DIV0, DIV5h, DIV3, DIV5, and DIV14 of retinoic acid differentiation. In the second approach we compared the extent to which the expression levels of select targets in differentiated H9c2 cells (DIV5 & DIV14) compare with those of primary cardiomyocytes. Refer to Table 3 for details of fold change results and statistical analyses.

**Figure 4.**
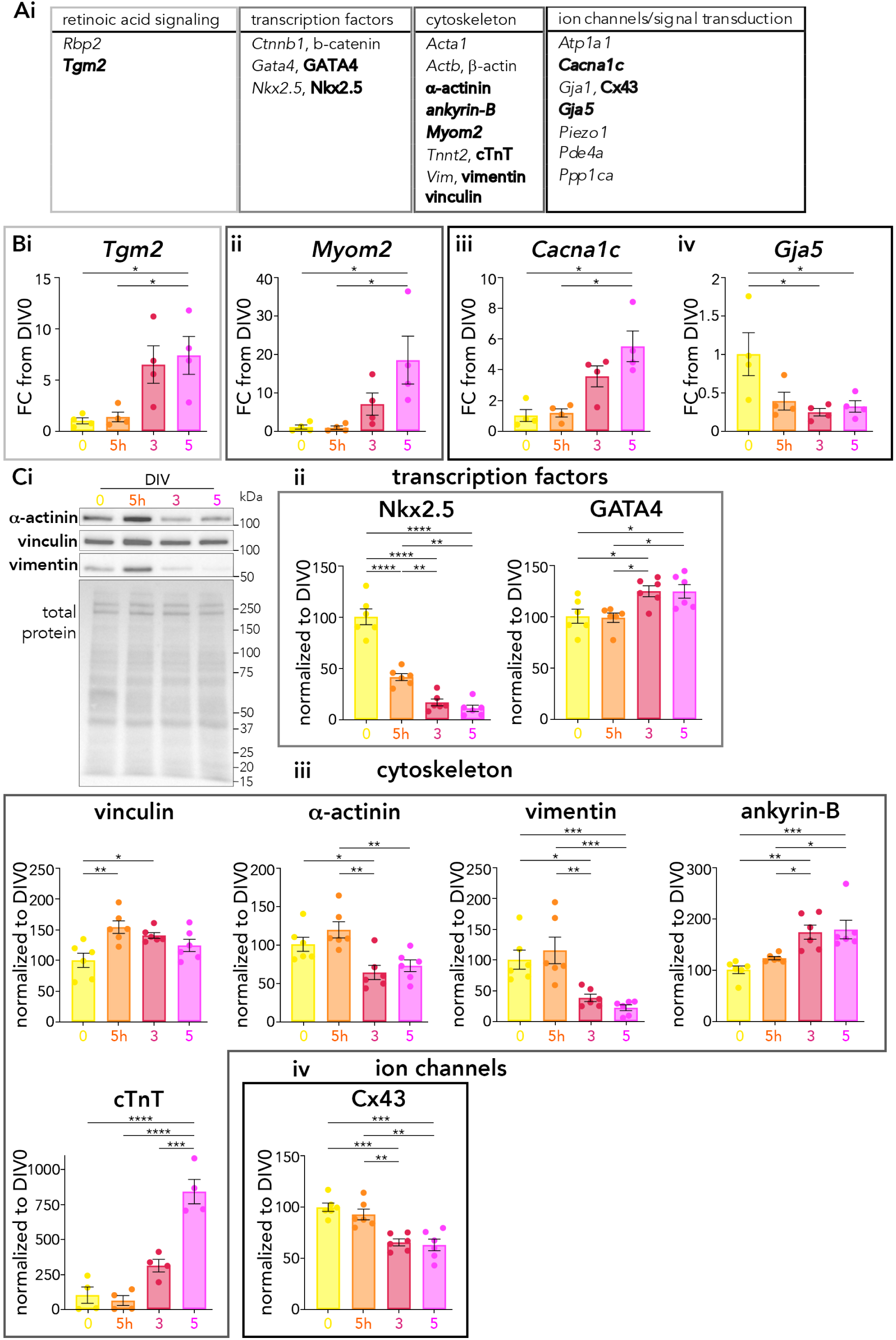
Differentiated H9c2 cells exhibit gene expression changes consistent with cardiomyocyte differentiation. (A) table of genes involved in retinoic acid signaling, transcription, cytoskeleton, ion flux regulation, and signal transduction were analyzed, bolded resulted in significant differences between groups. Transcript expression of (Bi) *Tgm2*, (Bii) *Myom2* and (Biii) *Cacna1c* increased during differentiation, while (Biv) *Gja5* decreased. (Ci) Representative western blot for some of the cytoskeletal proteins is shown. Protein expression of (Cii) transcription factor Nkx2.5 decreased while GATA 4 levels increased across differentiation. (Ciii) Cytoskeletal proteins levels increased and then returned to base line for vinculin, decreased for ⍺-actinin and vimentin, and increased for ankyrin-B and cTnT. Lastly, (Civ) the ion channel Cx43 levels decreased during differentiation. For RT-qPCR data: DIV0, DIV5h, DIV3, DIV5 (N=4 passages). For western blot data, DIV0, DIV5h, DIV3, DIV5 (N=6 passages, except cTnT N=4 passages). Data is presented as mean ± SEM. Results are graphed as Yellow = DIV0, Orange = DIV5h, Red = DIV3, Magenta = DIV5. ****, p < 0.0001; ***, p < 0.001; **, p < 0.01; *, p < 0.05; unlabeled, p > 0.05 by one-way ANOVA and post-hoc Tukey’s multiple comparison. Data is normalized to DIV0.

**Table 3.**
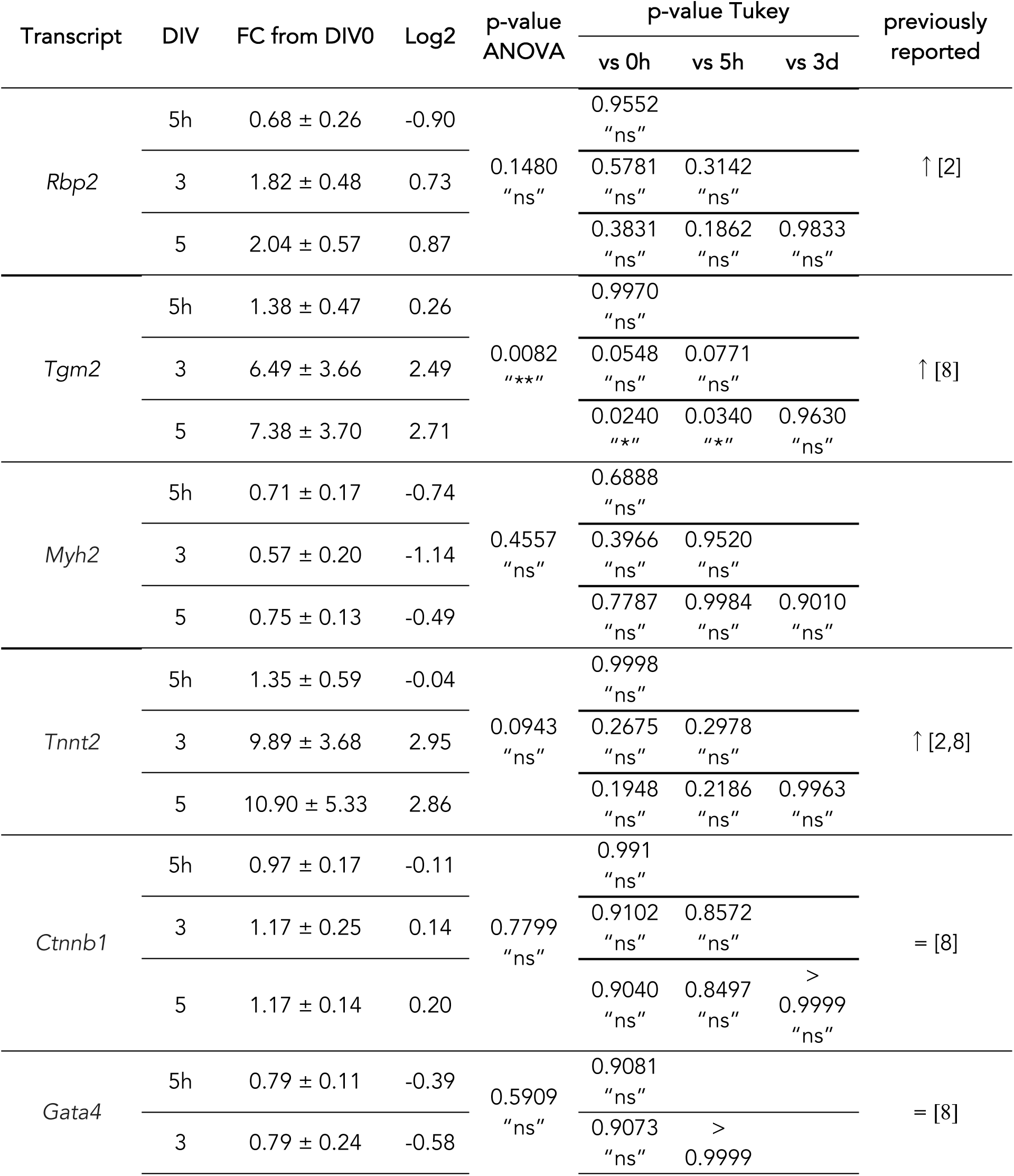
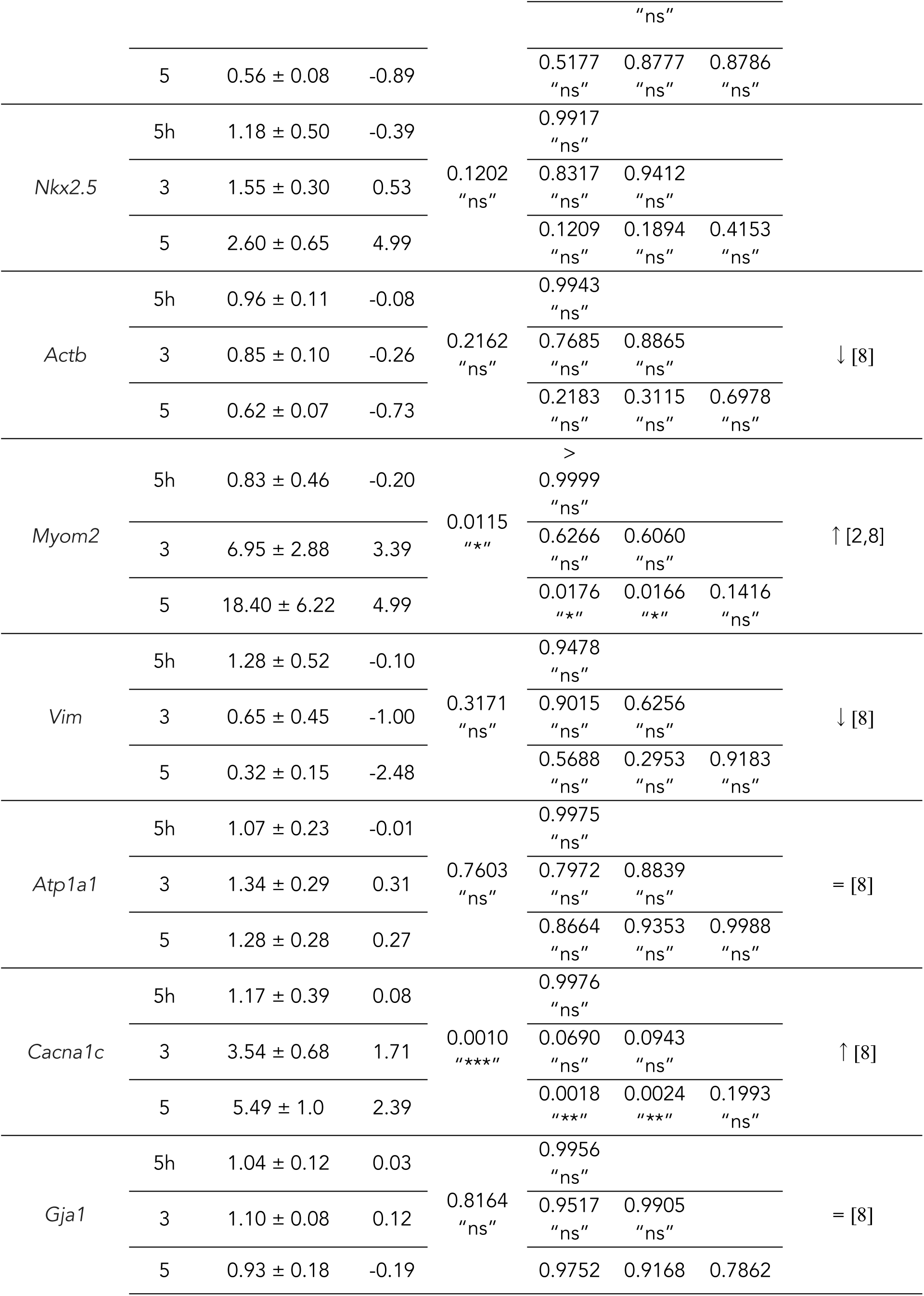
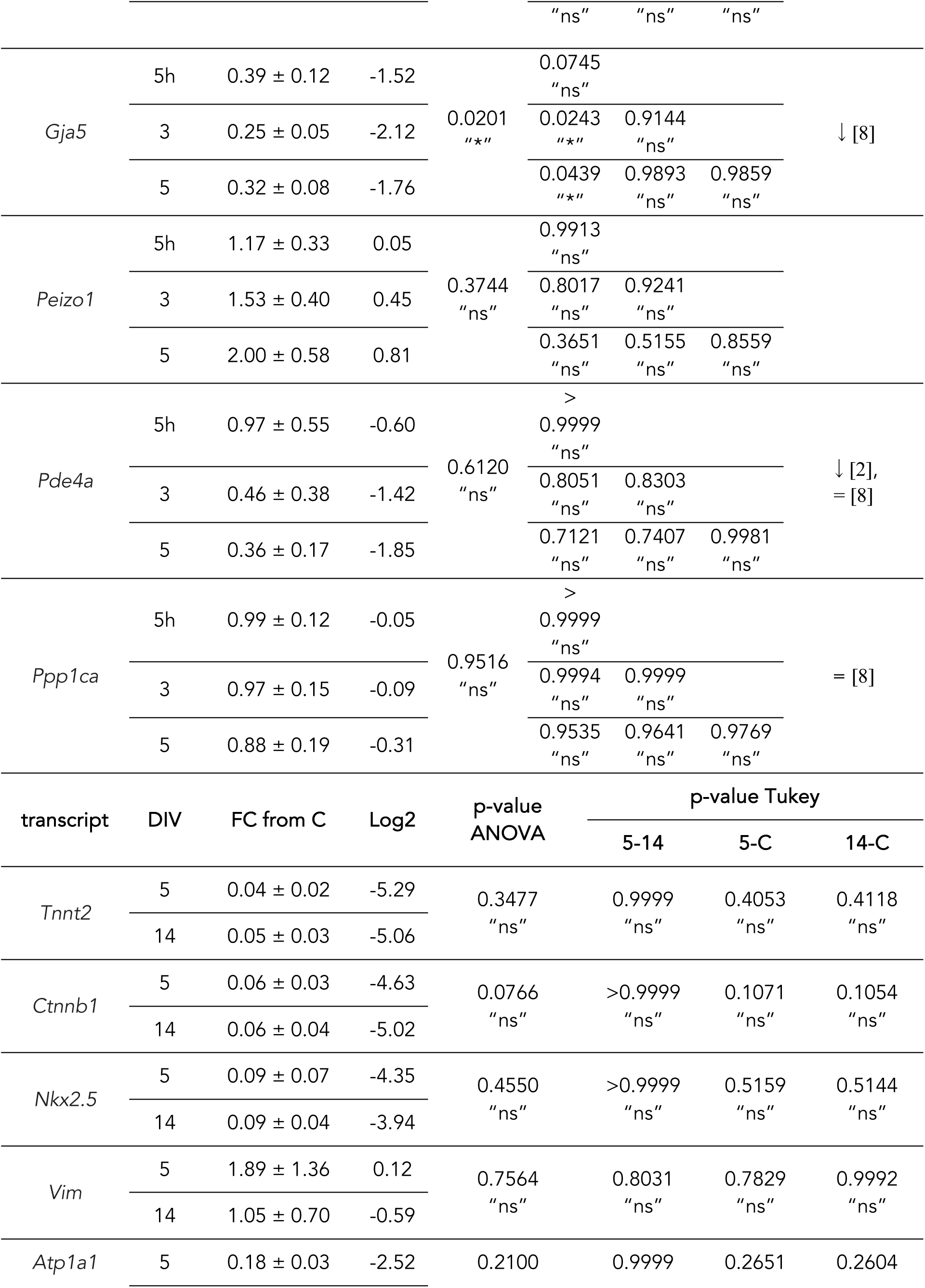
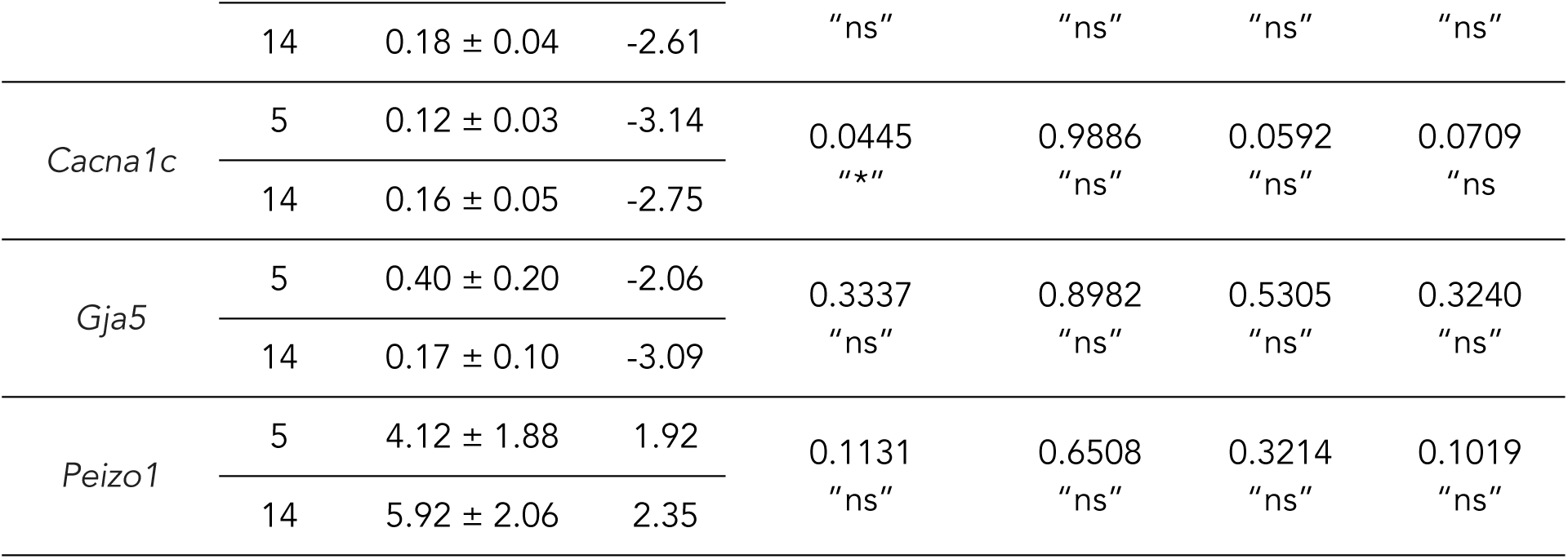
H9c2 cell transcript levels of key cardiomyocyte cytoskeletal proteins and ion channels increases over differentiation. For DIV5, which was compared to DIV0, and DIV5 and DIV14, which were compared to C, a one-way ANOVA was performed, if this provided a significant result, then the Tukey multiple comparisons test was used to determine which groups were significant. DIV0, DIV5h, DIV3, DIV5 (N=4 passages); DIV14, primary cardiomyocytes, P2 (N=3 passages, cultures, hearts); E15.5 (N=1 heart).

We confirmed retinoic acid signaling pathway activation, observing an increase in *Tgm2* (encodes for transglutaminase 2), at DIV5 compared to both DIV0 and DIV5h (Figure 4 Bi). Both *Myom2* (encodes for myomesin 2) and *Cacna1c* (encodes for L-type Ca^2+^ channel pore) also observed increased transcript levels, *Myom2* were elevated ∼18-fold ∼5 fold at DIV5 compared to both DIV0 and DIV5h, respectively (Figure 4 Bii, iii). *Gja5* (encodes for connexin43 (Cx43)) levels observed decreased levels at DIV3 and DIV5 (Figure 4 Biv)

We further evaluated protein expression changes for Cx43, as well as additional targets (Nkx2.5, GATA4, ⍺-actinin, Cx43, cTnT, vinculin, vimentin, β-actin, ankyrin-B) as shown in Figure 4 Ci, statistical analyses are in Table 4. Both transcription factors, Nkx2.5 and GATA4 levels were altered across differentiation (Figure 4 Cii). While Nkx2.5 levels decreased from DIV0 to DIV5, GATA4 levels increased. We also analyzed multiple cytoskeletal proteins (Figure 4 Ciii). Vinculin levels initially increased at DIV5h and DIV3. ⍺-actinin levels decreased at DIV3 and vimentin levels decreased at both DIV3 and DIV5. Ankyrin-B levels increased at DIV3 and DIV5. cTnT levels were increased at DIV5 as well. Lastly, Cx43 levels decreased during H9c2 cell differentiation at both DIV3 and DIV5.

**Table 4.** H9c2 cell protein levels of key cardiomyocyte cytoskeletal proteins and cardiac markers increases over differentiation. A one-way ANOVA was performed, if this provided a significant result, then the Tukey’s multiple comparisons test was used to determine which groups were significant. DIV0, DIV5h, DIV3, DIV5 (N=6 passages, except cTnT N=4 passages); DIV14, primary cardiomyocytes (C), P2, E15.5 (N=3 passages, cultures, hearts respectively).

| protein | DIV | normalized percent to DIV0 | p-value ANOVA | p-value Tukey |  |  | previously reported |
| --- | --- | --- | --- | --- | --- | --- | --- |
|  |  |  |  | vs 0 | vs 5h | vs 3 |  |
| Nkx2.5 | 5h | 41.3% ± 3.4 | <0.0001<br>"*****" | <0.0001<br>"*****" |  |  |  |
|  | 3 | 16.6% ± 3.3 |  | <0.0001<br>"*****" | 0.0078<br>"**" |  |  |
|  | 5 | 10.9% ± 3.1 |  | <0.0001<br>"*****" | 0.0011<br>"**" | 0.8297<br>"ns" |  |
| GATA4 | 5h | 98.5% ± 4.5 | 0.0043<br>"***" | 0.9979<br>"ns" |  |  | = [2] |
|  | 3 | 124% ± 5.3 |  | 0.0390<br>"**" | 0.0267<br>"**" |  |  |
|  | 5 | 124% ± 6.6 |  | 0.0409<br>"**" | 0.0280<br>"**" | >0.9999<br>"ns" |  |
| α-actinin | 5h | 119% ± 9.2 | 0.0013<br>"***" | 0.4883<br>"ns" | | | |
|  | 3 | 63.5% ± 10.5 |  | 0.0487<br>"**" | 0.0020<br>"***" |  |  |
|  | 5 | 72.3 ± 7.6 |  | 0.1760<br>"ns" | 0.0093<br>"***" | 0.9050<br>"ns" |  |
| Cx43 | 5h | 92.9% ± 5.3 | <0.0001<br>"*****" | 0.7108<br>"ns" |  |  |  |
|  | 3 | 65.6% ± 3.4 |  | 0.0002<br>"****" | 0.0028<br>"***" |  |  |
|  | 5 | 63.1% ± 5.7 |  | 0.0001<br>"****" | 0.0012<br>"***" | 0.9807<br>"ns" |  |
| cTnT | 5h | 61.3% ± 35.2 | <0.0001<br>"*****" | 0.9666<br>"ns" |  |  | ↑ [2,3] |
|  | 3 | 311% ± 44.9 |  | 0.1106<br>"ns" | 0.0510<br>"ns" |  |  |
|  | 5 | 839% ± 87.2 |  | <0.0001<br>"*****" | <0.0001<br>"*****" | 0.0002<br>"****" |  |
| vinculin | 5h | 154% ± 10.2 | 0.0043<br>"***" | 0.0033<br>"***" |  |  | ↓ [3,34] |
|  | 3 | 141% ± 4.58 |  | 0.0314<br>"**" | 0.7444<br>"ns" |  |  |
|  | 5 | 124% ± 10.0 |  | 0.2952 | 0.1522 | 0.6297 |  |

|  |  |  |  | "ns" | "ns" | "ns" |
| --- | --- | --- | --- | --- | --- | --- |
| vimentin | 5h | 115% ± 21.5 | 0.0002<br>"****" | 0.8661<br>"ns" |  |  |
|  | 3 | 38.0% ± 6.0 |  | 0.0224<br>"★" | 0.0040<br>"★★" |  |
|  | 5 | 21.9% ± 4.8 |  | 0.0036<br>"★★" | 0.0006<br>"****" | 0.8425<br>"ns" |
| β-actin | 5h | 131% ± 14.0 | 0.0663<br>"ns" | 0.4412<br>"ns" |  |  |
|  | 3 | 149% ± 11.9 |  | 0.1076<br>"ns" | 0.8139<br>"ns" |  |
|  | 5 | 153% ± 13.1 |  | 0.0756<br>"ns" | 0.7132<br>"ns" | 0.9977<br>"ns" |
| ankyrin-B | 5h | 122.5 ± 3.1 | 0.0003<br>"****" | 0.5575<br>"ns" |  |  |
|  | 3 | 172.7 ± 13.7 |  | 0.0018<br>"★★" | 0.0352<br>"★" |  |
|  | 5 | 177.9 ± 17.9 |  | 0.0009<br>"****" | 0.0182<br>"★" | 0.9896<br>"ns" |
| protein | DIV | normalized<br>percent to C | p-value<br>ANOVA | p-value Tukey |  |  |
|  |  |  |  | 5-14 | 5-C | 14-C |
| Nkx2.5 | 5 | 10.3% ± 5.7 | 0.0005<br>"****" | 0.9758<br>"ns" | 0.0009<br>"****" | 0,0010<br>"★★" |
|  | 14 | 12.9% ± 8.9 |  |  |  |  |
| α-actinin | 5 | 169% ± 10.7 | 0.0085<br>"★★" | 0.0436<br>"★" | 0.0075<br>"★★" | 0.3271<br>"ns" |
|  | 14 | 123% ± 4.0 |  |  |  |  |
| Cx43 | 5 | 41.5% ± 19.1 | 0.3558<br>"ns" | 0.9895<br>"ns" | 0.4522<br>"ns" | 0.3878<br>"ns" |
|  | 14 | 35.2% ± 10.6 |  |  |  |  |
| cTnT | 5 | 11.9 % ± 9.2 | 0.0013<br>"★★" | 0.9963<br>"ns" | 0.0023<br>"★★" | 0.0022<br>"ns" |
|  | 14 | 10.7% ± 5.9 |  |  |  |  |
| vinculin | 5 | 205% ± 4.5 | 0.0352<br>"★" | 0.6801<br>"ns" | 0.0344<br>"★" | 0.1005<br>"ns" |
|  | 14 | 178% ± 11.0 |  |  |  |  |
| vimentin | 5 | 725% ± 42.0 | <0.0001<br>"*****" | <0.0001<br>"*****" | <0.0001<br>"*****" | 0.9229<br>"ns" |

Next, to determine which differentiation timepoint best approximates the primary cardiomyocyte phenotype at a gene expression level, we compared the levels of select targets from DIV5 or DIV14 differentiated H9c2 cells to those in cardiomyocytes (Figure 5 A). There were no significant differences between DIV5, DIV14, or primary cardiomyocyte transcript expression for any of the genes we analyzed (see Figure 5 A for the list of genes). For analysis of protein levels, we included embryonic (E15.5) and early postnatal (P2) mouse heart tissue samples as positive controls (Figure 5 Bi). Vinculin, ⍺-actinin, and vimentin levels at DIV5 but not DIV14 were significantly different from primary cardiomyocytes (Figure 5 Bii). While the levels of cTnT and Nkx2.5 at both DIV5 and DIV14 were significantly reduced in H9c2 cells compared to primary cardiomyocyte levels (Figure 5 Bii, Biii)

**Figure 5.**
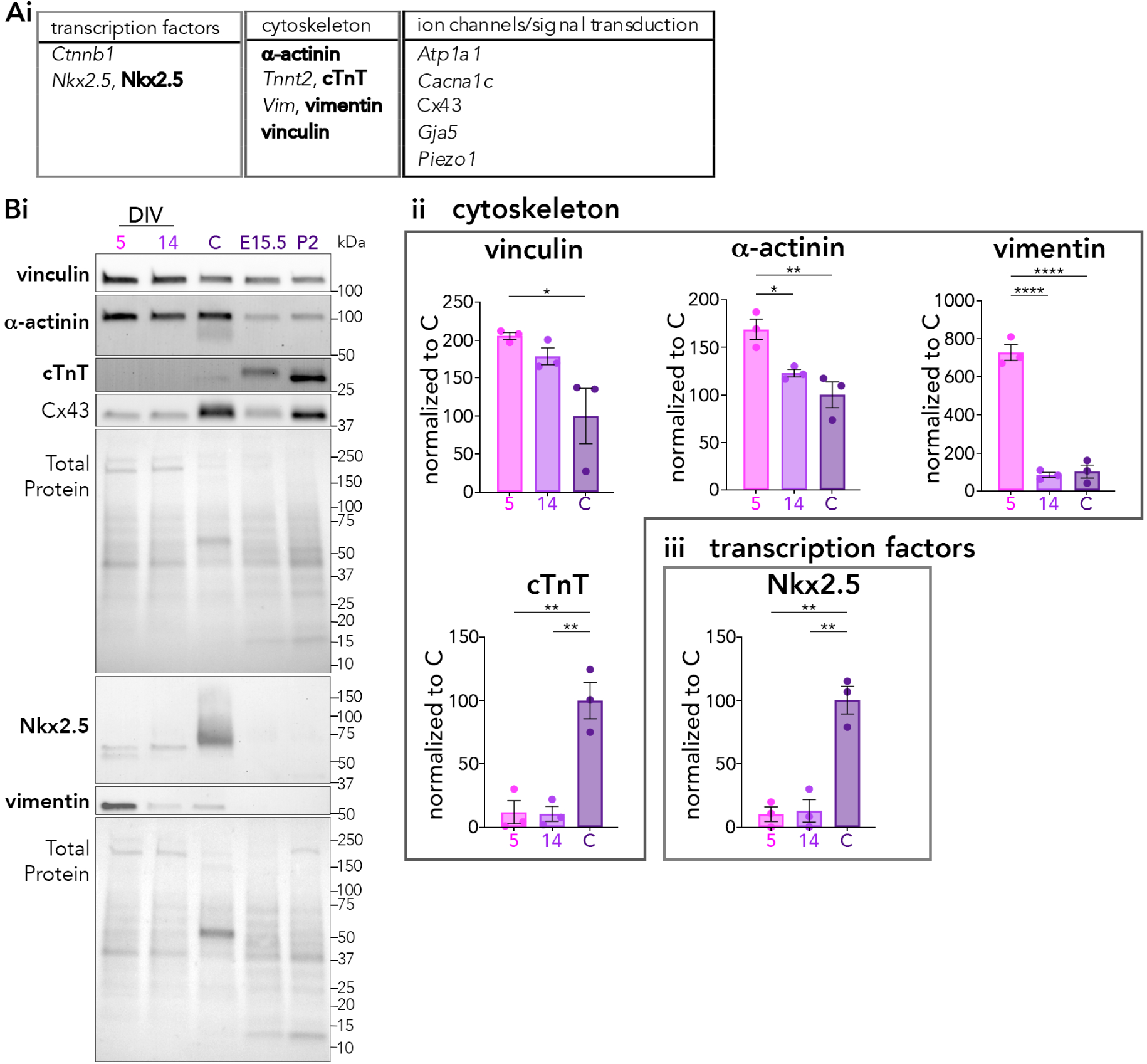
Differentiated H9c2 cells at DIV14 more closely resemble primary cardiomyocytes than DIV5. (A) table of genes involved in retinoic acid signaling, transcription, cytoskeleton, ion flux regulation, and signal transduction were analyzed, bolded resulted in significant differences between groups. (Bi) Representative western blot for some of the cytoskeletal proteins is shown. (Bii) Protein expression of several cytoskeletal proteins were significantly different at DIV5 but not at DIV14 including vinculin, ⍺-actinin, and vimentin, while cTnT levels were significantly decreased at both DIV5 and DIV14 compared to primary cardiomyocytes. (Biii) Similarly, the levels of transcription factor Nkx2.5 were also significantly decreased at both DIV5 and DIV14 compared to primary cardiomyocytes. For western blot data DIV5, DIV14 (N = 3 passages; primary cardiomyocytes (C) (N = 3 cultures); P2, E15.5 (N=3 hearts). Data is presented as mean ± SEM. Results are graphed as Yellow = DIV0, Orange = DIV5h, Red = DIV3, Magenta = DIV5. ****, p < 0.0001; ***, p < 0.001; **, p < 0.01; *, p < 0.05; unlabeled, p > 0.05 by one-way ANOVA and post-hoc Tukey’s multiple comparison. Data is normalized to primary cardiomyocytes (C).

Together, these data are in line with many previously reported changes in the expression of several key ion channels and cytoskeleton-associated genes with H9c2 cell differentiation alters, and identify several new ones, including ⍺-actinin and ankyrin-B. Overall, these results suggest DIV14 cells most closely resemble the primary cardiomyocyte gene expression.

### Summary of major changes and associated experimental recommendations

The use of H9c2 cells across different timepoints, DIV0-3, DIV5, and DIV14, may be useful for different studies depending on the research question. H9c2 cells at DIV0 represent the undifferentiated baseline, with smaller, mononucleated cells lacking calcium signaling and with decreased expression of several key cardiomyocyte genes (e.g., *Myom2, Cacna1c*, cTnT). For detecting early changes to gene expression timepoints before DIV5 may be of use (such as DIV5h or DIV3). DIV5 cells highlight initial changes to cell eccentricity and trends towards cell shape changes, the emergence of spontaneous Ca²⁺ transients, and gene expression changes associated with mature cardiomyocytes. By DIV14, cells exhibit pronounced cardiomyocyte-like features, such as multinucleation, actin clusters, Ca²⁺ handling, and gene profiles more closely resembling primary cardiomyocytes. H9c2 cells at DIV14 are likely the best candidate for morphological and functional outputs while DIV5 and earlier cells are beneficial for studying gene expression changes preceding the morphological changes.

## Discussion

Here we conducted a parallel analysis of H9c2 cell morphology, spontaneous Ca^2+^ handling, and gene expression over retinoic acid differentiation. We found that differentiated H9c2 cells recapitulate specific aspects of cardiomyocyte differentiation in all three areas, as summarized in Figure 1. This information will facilitate the optimal use of differentiated H9c2 cells to advance understanding of the mechanisms underlying cardiomyocyte differentiation with health and disease applications.

Our quantitative analysis of H9c2 cell morphology revealed marked changes in multiple aspects across H9c2 cell differentiation. These include increases in mean area, eccentricity, and multinucleation (Figure 2). The most robust morphological differences were observed at DIV14, expanding upon the findings observed by others [3,8] by incorporating additional measures including quantifying length, perimeter, and eccentricity in addition to area and multinucleation. These findings suggest that DIV14 is the best model for investigating elements related to morphology and function. Early gene expression changes related to cytoskeletal organization. Such as upregulation increased expression of *Cacna1c*, which encodes for the L-type Ca^2+^ channel pore, not only initiates Ca²⁺ signaling but also supports downstream calcium-dependent cytoskeletal remodeling pathways. Modulation of proteins like vinculin, α-actinin, vimentin, and ankyrin-B, likely contribute to the increases in cell length, area, and eccentricity observed across differentiation. These structural changes are further accompanied by the emergence of actin clusters, which became significantly more prevalent at DIV14, as revealed by super-resolution microscopy (Figure 2 D,E). Changes in actin are key for the development of cardiomyocyte cell shape and sarcomere formation [21–23] and while the role of actin clusters has not been thoroughly characterized in cardiomyocytes, they are known to play important roles in regulating actin dynamics in other cell types [24–26]. Therefore, differentiated H9c2 cells could provide a model in which to investigate the generation and role of actin clusters in cardiomyocyte development. Spontaneous Ca²⁺ transients first appeared at DIV5 (Figure 3), aligning with early increases in calcium-handling gene expression, such as *Cacna1c*. Although average Ca²⁺ dynamics in H9c2 cells did not reach primary cardiomyocyte levels, distribution shifts at DIV14 more closely resemble the dynamics of primary cardiomyocytes compared to DIV5. The morphological changes, presence of spontaneous Ca²⁺ transients, and gene expression changes at DIV14 the most suitable to model cardiomyocytes.

### Translational implications

The ability of differentiated H9c2 cells to model key morphological, functional, and gene expression features of cardiomyocytes highlights their relevance as a model system for investigating the cellular and molecular mechanisms underlying cardiac development and disease. While this differentiation is nuanced, the ability to model cardiomyocyte differentiation in a robust and easily manipulated cell line allows for experiments including prioritizing hypotheses before advancing to more complex models and high throughput screenings of therapeutics. Furthermore, the ease of transfection and use of inducible vectors enables the investigation of the impacts of genetic variants at various stages of cardiomyocyte differentiation (Figure S2). Overall, the morphological, functional, and gene expression changes that occur across differentiation indicate that H9c2 cells are useful for identifying disease mechanisms, therapeutic targets, and developing interventions.

### Limitations

One of the challenges in measuring the morphology of H9c2 cells is the proximity of cells; however, this feature itself is a critical process during cardiomyocyte development and in diseased states [27,28]. As a result, the segmentation was challenging given the strong expression of actin stress fibers throughout the cell. To measure the morphology of H9c2 cells we identified the number of segmentable cells (cells with clear boundaries) in an FOV and randomly selected 3 cells. Inherently, this may have favoured the more undifferentiated cells in the FOV as they are less aligned by nature. Future studies could employ cell perimeter stains such as wheat germ agglutinin for easier analysis and to better quantify the whole region of interest.

Due to logistical limitations, we performed the Ca^2+^ imaging for different experimental groups on different days. In future work, we suggest that a final stimulated Ca^2+^ spike could be used to normalize amplitudes, for example.

In the present study, for the primary cardiomyocytes, we performed mixed sex cultures. Given the known sex differences in heart disease, the development of single heart cultures may explain some differences between H9c2 cells and the models as H9c2 cells were isolated from a female left ventricle. Furthermore, the quantification of primary cardiomyocyte culture composition may also explain differences if some cultures contain more endothelial or fibroblast cells. And given the benefit of having a culture more resembling what is seen *in vivo* it may be worthwhile to maintain the presence of endothelial cells and fibroblasts. Finally, in future work, the sex of the animal from which all heart tissues are derived should be recorded.

### Conclusions

The H9c2 rat ventricular cardiomyoblast cell line has been used to model cardiomyocyte development and disease. Here we quantified changes in three broad areas of H9c2 cell biology across five and fourteen days of retinoic acid differentiation: morphology, Ca²⁺handling, and gene expression. Our quantitative assessment of shape, spontaneous Ca²⁺handling, and gene expression (both early at DIV5h and DIV3 as well as later at DIV14) worked to address some of the knowledge gaps in the field to fully facilitate the use of these cells. Depending on the context, we compared to DIV5 cells or primary cardiomyocytes. Overall, we saw differentiation-induced changes in all three areas.

Increased cell size, multinucleation, development of spontaneous Ca^2+^ transients, and the regulation of several key genes involved in these processes position H9c2 cells as a high-throughput model for studying genetic and environmental factors affecting cardiomyocyte biology, with the advantage of easier and shorter culturing protocols. Additionally, their ease of transfection allows for targeted investigation of specific genes and pathways, and the impacts of variants on cardiomyocyte differentiation. Our comprehensive characterization indicates that H9c2 cells are a useful model in investigating components of cardiomyocyte differentiation and may be particularly useful for understanding disease mechanisms and for screening potential therapeutic interventions. However, the heterogenicity of differentiated H9c2 cell analysis should take into account the spread of differentiated cells and the addition of other models such as primary cardiomyocytes and iPSC-derived cardiomyocytes may be of value to validate the findings.

## Materials and Methods

All data that support the findings of this study are available from corresponding authors upon reasonable request.

### Cell culture

The H9c2 rat ventricular-derived cardiomyoblast cell line was obtained from the American Type Culture Collection (ATCC, CRL-1446). H9c2 cells were cultured in Dulbecco’s Modified Eagle Medium (DMEM, ATCC, 30-2002) supplemented with 10% fetal bovine serum (FBS) (Thermo Fisher, 12483020, reserve 10498874), 100 U/mL penicillin, and 100 μg/mL streptomycin (Thermo Fisher, 15140122) at 37°C in 5% CO_2_. Cells were kept at or below 15 passages and passaged at 70-80% confluence to prevent the loss of differential potential. Each vial was tested for mycoplasma contamination using the universal mycoplasma detection kit (ATCC, 30-1012K).

H9c2 cells were differentiated as previously described [1]. Briefly, cells to be collected within 5 days were seeded at 8,333 cells/cm^2^ for RT-qPCR, Western blotting, and live cell Ca^2+^ imaging experiments, and 8,000 cells/cm^2^ for immunocytochemistry experiments. For cells collected at 14 days, cell densities were decreased by half to prevent cells from becoming confluent and detaching. One day after seeding, FBS was decreased from 10% to 1% and media was supplemented with 1 µM all-trans retinoic acid (ATRA) (Sigma, R2625-50MG). ATRA was added daily, in the dark, and cells were collected prior to ATRA treatment (DIV0), at 5 hours (DIV5h), 3 days (DIV3), 5 days (DIV5), and 14 days (DIV14) post ATRA treatment. Sample size was chosen based from the resource equation [29].

One day after seeding, cells were transfected with a tetracycline inducible pCMB6-AC-GFP-rtTA plasmid (Origene, PS100125) using jetPEI (Figure S2). 4 µL of jetPEI diluted in 150 mM NaCl was added to 2 µg of plasmid DNA also diluted in 150 mM NaCl, the mixture was incubated for 15 minutes, and then added dropwise to the cells in DMEM containing no antibiotics. One day after transfection, media was changed to DMEM containing 1% FBS, 100 U/mL penicillin, and 100 μg/mL streptomycin, and was treated daily with 1 µM ATRA. To induce GFP expression, 10 ng/mL Doxycycline hyclate (Sigma, D9891-1G) was added to the differentiation media.

### Experimental Animals

All animal procedures were approved by the University of Victoria Animal Care Committee and performed in accordance with the guidelines set by the Canadian Council on Animal Care. Male and female embryonic day (E)15.5– postnatal day (P)2 mice were used in this study. C57BL/6J mice were obtained from The Jackson Laboratory (#000664). Mice were housed under a 12/12 hour light/dark cycle starting at 8 A.M., with food and water ad libitum; temperature was maintained between 20°C and 25°C and humidity at 40– 65%. Sample size was chosen based from the resource equation [29].

### Heart Tissue Extraction

E15.5 and P2 heart tissue was flash frozen in liquid nitrogen and stored at −80°C. E15.5 heart tissue intended for RNA analysis was stored in RNAlater Stabilization Solution (Thermo Fisher, AM7020). Heart tissue was homogenized in a Dounce homogenizer on ice with either 100 µL TRIzol (Thermo Fisher, 15596026) per 5-10 mg tissue (RNA) or RIPA lysis buffer (9.1 mM PBS, 150 mM NaCl, 1.7 mM monobasic sodium phosphate, 9.1 mmol/L dibasic sodium phosphate, 1% Igepal, 0.5% w/v sodium deoxycholate, 0.1% sodium dodecyl sulphate (SDS), 10 µL/mL protease inhibitor cocktail (Sigma, P8340-5ML)) (protein). RNA was extracted from tissue samples after incubating on ice for 5 minutes and then stored in TRIzol at −80°C. Protein lysates were pushed through a 27 ½ gauge needle 2 times and incubated on ice for 30 minutes; samples were then centrifuged at 13500 x g for 20 minutes at 4°C and the supernatant was also frozen at −80°C.

### Primary culture isolation

Mouse cardiomyocytes were isolated according to standard methods previously described [30]. 6-10 hearts were used per each given culture, representing 1 biological replicate (N). Briefly, hearts were extracted from P1-P2 mice and transferred into phosphate-buffered saline solution (without Ca^2+^ or Mg^2+^) containing 20 mM 2,3-Butanedione monoxime (BDM) (Sigma, B0753). In a sterile cell culture hood, the hearts were washed, minced, and submerged in isolation medium consisting of Hanks’ Balanced Salt Solution (HBSS) with 20 mM BDM and 0.025% Trypsin (Gibco, 15090-046) for incubation overnight under gentle agitation at 4°C. After incubation, the isolation medium was removed, and the tissue was oxygenated in Leibovitz’s L-15 medium containing 10 mM BDM and 0.75 mg/ml of Collagenase/Dispase® mixture (Roche, 10269638001), then placed in a shaking incubator for 30 minutes at 37°C. To release cells into suspension, the tissue fragments were gently triturated and filtered through a 100 µm nylon mesh cell strainer.

Cells were centrifugated at 200 x g for 5 minutes to remove fibroblasts and endothelial cells. The resulting pellet was resuspended in Dulbecco’s modified Eagle’s medium (DMEM) supplemented with 5% Heat Inactivated Horse Serum (Thermo Fisher, 26050070) and plated in a cell culture dish (a pre-plating step that removes many of the remaining fibroblasts and endothelial cells). After a 2-3 hour incubation, non-adherent cardiomyocytes were collected and plated on pre-coated collagen coverslips (Neuvitro, NEU-GG-12-1.5-COLLAGEN) or onto collagen from calf skin (Fisher, ICN16008410) coated 35 mm glass bottom imaging dishes at a density of 1.5 x 10^5^ cells per cm^2^. The following day, plating medium was replaced by DMEM containing 5% HIHS, 100 U/mL penicillin, and 100 μg/mL streptomycin for maintenance.

### Immunocytochemistry

H9c2 cells were seeded onto fibronectin coated coverslips either purchased from Neuvitro (NEU-GG-12-1.5-FIBRONECTIN) or coated in lab using fibronectin bovine plasma (Sigma, F1141-1MG). Coverslips were fixed with 4% paraformaldehyde for 5 minutes at DIV0, DIV5, and DIV14, followed by three washes with PBS containing Ca^2+^ or Mg^2+^. Cells were permeabilized with 0.1% Tween-20 for 5 minutes, washed three times in PBS as before, blocked with 1% BSA and 22.52 mg/mL glycine in PBS-T, and then were incubated with Hoechst (1:500) and phalloidin (1:500; 555, Invitrogen, A34055; Acti-Stain 670, Cytoskeleton, PHDN1) diluted in 1% BSA in PBS overnight at room temperature for 1 hour. Following secondary incubation, coverslips were washed three additional times, and mounted with ProLong Gold Antifade Mountant (Thermo Fisher, P10144).

For transfected cells, fixation with 4% paraformaldehyde was performed as previously described, followed by three washes in dPBS and mounting with ProLong Gold Antifade Mountant.

### Microscopy

Cells were imaged using a Leica TCS SP8 confocal microscope with HyD detectors. For DIV0 versus DIV5 comparisons, imaging was performed using a 40X oil-immersion objective set at 0.75 zoom (1.30 NA, pinhole = 65.28 µm, 3904 x 3904 (pixel size = 74.46 x 74.46 nm)). For DIV5 versus DIV14 cells, images were acquired using the 20X dry-objective (0.7 NA, pinhole = 60.69 µm, 3800 x 3800 (pixel size =148.92 nm x 148.92 nm)).

For transfected cells, images were acquired using the 40X oil-immersion objective set at 0.75 zoom (1.30 NA, pinhole = 65.08 µm, 2048 x 2048 (pixel size = 141.98 nm x 141.98 nm)). GFP was excited using the 488 nm Argon laser at 10% intensity.

STED images were acquired using a 100X oil-immersion objective (1.40 NA, pinhole = 117.5 µm, pixel size optimized to 20 µm x 20 µm) at a zoom of 2. Phalloidin 555 was excited using the 514 Argon laser at 20% intensity. Fluorescence was captured using the HyD detector using the 660 STED laser at 25% 3D, 80% intensity and 70-90% power.

For all images presented, brightness and contrast were uniformly adjusted for presentation purposes.

### Morphology Analysis

To analyze morphological features of H9c2 cells, a custom Matlab script was created: Morphology Analysis Overview A (code available at https://github.com/SwayneLab/Quantification-of-H9c2-cell-differentiation-/). Images were blinded before analysis. First, the analysis region for each cell was marked by manually drawing a polygon around it. The script processed a binarized image where cell pixels are set to one and background pixels to zero. Zeros were padded around the image to ensure accurate boundary detection at the edges. The cell boundary was identified by examining each pixel and its eight surrounding neighbours in a three x three grid, with boundary pixels identified whenever a surrounding pixel was zero. Next, a covariance matrix was created from these repositioned coordinates. An eigenvalue decomposition was performed on this matrix to find the principal directions in which the cell extended. These principal directions correspond to the longest (major) and shortest (minor) dimensions of an ellipse that could fit around the cell.

Length, area, and eccentricity were calculated by first finding the center point, or centroid, of each cell, averaging the positions of all the pixels that made up the cell. This average point helped reposition the cell’s pixel distribution to the center of the coordinate system for easier analysis. Length was calculated from the square root of the largest eigenvalue (major axis) and eccentricity was calculated from the ratio of largest eigenvalue (major axis) to smallest eigenvalue (minor axis). The perimeter was calculated by counting the number of boundary pixels, and area was calculated from filling in pixel area between the boundary pixels. The number of nuclei per analyzed cell was calculated manually. All plots and statistics were performed in Prism 10 for Mac OS X (version 10.3.1, GraphPad Software, Inc.).

### Calcium imaging and analysis

Isolated mouse cardiomyocytes seeded in collagen-coated 35 mm glass bottom imaging dishes (ibidi, 81218-200) were loaded with cell-permeant 4 µM Fluo-4 AM (Invitrogen, F-14201) for 35 minutes. H9c2 cells seeded in fibronectin-coated imaging dishes (ibidi, 81218-200) were loaded with 1.5 µM Fluo-4 AM at room temperature for 25 minutes.

Incubation time, temperature, and loading concentrations were optimized for each cell type to prevent dye toxicity and compartmentalization. The cells were subsequently washed using HBSS with 10 mM HEPES (Stem Cell Technologies, 37150) and incubated for 20 minutes to allow for de-esterification of the AM esters. All samples were imaged in complete FluoroBrite™ DMEM media (Gibco™, A1896701) at 37°C and 5% CO_2_ in a Tokai Hit Stage Top Incubation System mounted on the Leica SP8. Fluo-4 was excited with the 488 nm argon laser line at 20% intensity through a 20x objective (0.70 NA, pinhole =242.65 µm). Using the Leica SP8 resonant scanner, cardiomyocytes were imaged at 27.7 FPS for 1 minute and H9c2 cells were imaged at 14.6 FPS for 2 minutes. A minimum of 2 dishes were imaged per biological replicate with 2-5 fields of view captured per dish.

Imaging files were initially processed by EZcalcium, an open-source, automated data analysis package [31]. Part of the Matlab-based toolbox, the automated ROI detection module was run first to detect all active cells, followed by the ROI refinement module to semi-automatically eliminate dying cells and those with a poor signal/noise. Minor movement artefacts in some FOV were not deemed significant by the motion correction module. For quantification of the extracted data files, a series of Matlab scripts based on those published [20] were employed. All data were analyzed and plotted in Prism 10 for Mac OS X (version 10.3.1, GraphPad Software, Inc.), frequency distributions were created in R (version 4.4.0).

### RNA extraction

RNA was isolated using TRIzol as in [32]. Briefly, RNA from H9c2 cells or primary cardiomyocytes was collected by adding 0.4 mL TRIzol per 1×10^5^-1×10^7^ cells. Extracted RNA was stored at −80°C.

RNA integrity was checked using the Agilent RNA 6000 Nano Kit (Agilent, 5067-1511). Briefly, to minimize secondary structure, 2 µL RNA was denatured at 70°C for 2 min.

1 µL was loaded into the primed chip containing the gel-dye mix and RNA 6000 nano marker. Once loaded, the chip was vortexed at 2400 rpm for 60 sec in the IKA vortex mixer, loaded into the Agilent 2100 bioanalyzer, and the total RNA (eukaryotic) run was started within 5 minutes. The RNA integrity number (RIN) scores were above 8.8 for H9c2 samples, 7.6 for P2 heart tissue and primary cardiomyocytes, and 1.5 for the E15.5 sample which showed both the 18S and 28S bands.

### RT-qPCR

1000 ng of RNA underwent treatment with DNase1 (Thermo Fisher, EN0521) to digest any potential double-stranded DNA isolated with the RNA. cDNA was synthesized from 500 ng of the DNase1 treated RNA using the Maxima First Strand cDNA Synthesis Kit using random primers (Thermo Fisher, K1652). For each RT+ reaction, an RT-reaction was also run. A GAPDH PCR reaction was run to confirm the cDNA synthesis and validate the RT-controls on a 2% agarose gel. qPCR reactions for genes of interest were performed on a QIAquant 96 well 5 plex qPCR machine (Qiagen, 9003010). Using PrimeTime qPCR probe assays (IDT) and PrimeTime gene expression master mix (IDT, 1055772), 100 µg of RNA was run in 10 µL reactions. The probe assays used are listed in Table S2. Standard cycling conditions were run: polymerase activation at 95°C for 3 minutes, followed by 45 cycles of denaturation at 95°C for 15 seconds and annealing/extension at 60°C for 1 minute. RT-qPCR data was inputted into geNorm to determine the best normalization strategy [33]. To normalize target genes, relative expression was calculated using the geometric mean of two housekeeping genes, *Polr2a* and *Ppia*. Graphs were created in Prism 10 for Mac OS X (version 10.3.1, GraphPad Software, Inc.).

### Western Blotting

Adhered cells were washed 2 times in dPBS and lysed in RIPA lysis buffer (9.1 mM PBS, 150 mM NaCl, 1.7 mM monobasic sodium phosphate, 9.1 mmol/L dibasic sodium phosphate, 1% Igepal, 0.5% w/v sodium deoxycholate, 0.1% sodium dodecyl sulphate (SDS), 10 µL/mL protease inhibitor cocktail (Sigma, P8340-5ML)) by scraping cells off the plate or coverslip on ice. Cell lysates were then treated as described for the mouse heart tissue. Protein content was determined using the DC protein assay (Bio-rad, 5000112). 10 µg of protein was incubated in Laemmli sample buffer (1X: 0.08 mM bromophenol blue, 0.05 M Tris-HCl pH 6.8, 0.058 M SDS, 3.33% v/v glycerol) at room temperature under reducing conditions (5% dithiothreitol (DTT) and 5% β-mercaptoethanol), separated by SDS-PAGE on TGX-stain free gels (Bio-rad, 4561094) and transferred to either a 0.2 μm pore PVDF membrane (Bio-rad, 1620177) for peroxidase detection using the Syngene G:Box imager or a 0.45 µm pore LF-PVDF membrane (Sigma, IPFL00010) for fluorescent detection using the Licor Odyssey CLx Imager. For Western blots using the peroxidase-based detection, the membrane was blocked with 5% fat free instant skim milk in TBS-T, then incubated with indicated antibodies in the blocking solution, and protein bands were visualized using the Clarity Western enhanced chemiluminescence substrate (Bio-Rad, 1705061). For Western blots using the fluorescent-based detection, the membrane was blocked with intercept (TBS) blocking buffer (Li-cor, 927-60003), incubated with indicated antibodies in the blocking solution containing 0.1% Tween 20 and 0.01% SDS (secondary only). All proteins were normalized to total protein using Ponceau S Solution (0.1% w/v in 5% acetic acid) (Sigma, P7170-1L) or the Revert 700 Total Protein Stain (Li-cor, 926-11010) and quantified with ImageJ. The antibodies and concentrations used are listed in Table S3. Graphs were created in Prism 10 for Mac OS X (version 10.3.1, GraphPad Software, Inc.).

## Acknowledgements

This work was supported by a Canadian Institutes for Health Research Project Grant (CIHR; PJT-169064) awarded to LTA and LAS as well as from CIHR grants (PJT-185887, PJT-189953) awarded to LAS. NY was supported by a Vanier Canada Graduate Scholarship (Canadian Institutes of Health Research Vanier) and University of Victoria Donor Awards. We acknowledge that this research was conducted on the traditional territory of the Ləkʷ̓əŋən (Songhees and Esquimalt) Peoples, and the Ləkʷ̓əŋən and W̱SÁNEĆ Peoples whose historical relationships with the land continue to this day. We also acknowledge the Gitxsan peoples and their leadership as this work was initiated for and is foundational for our ongoing research partnership on the mechanisms of inherited cardiac disease. The progress of this work has been presented at in-person community gatherings, and virtually by NY.

## Author contributions

NY, JR, LA, and LAS conceived and designed the experiments. NY conducted the cell culture work, morphological imaging, and biochemical assays, as well as advised in the morphological and Ca^2+^ analyses. MRM created the morphological analysis, which both MRM and RS conducted, NY then graphed and analyzed. JR and NY isolated primary cardiomyocyte cultures. JR performed all Ca^2+^ imaging and analysis, as well as directed the creation and statistical analysis for Figure 3. NY conducted all statistical analysis. NY and KWLF created all figures. NY, JR, RS, KWLF, LEWS, LA, and LAS wrote and edited the manuscript.

[TAB]

## Abbreviations

iPSC: induced pluripotent stem cell
Ca_v_1.2: protein pore subunit of the of the L-type calcium channel encoded by the *CACNA1C* gene
Ca^2+^: Calcium ions
[Ca^2+^]i: Internal calcium ion concentration
STED: Stimulated emission depletion
NaCl: Sodium chloride
E15.5: Embryonic Day 15.5
P2: Postnatal Day 2
DMEM: Dulbecco’s Modified Eagle Medium
FBS: Fetal bovine serum
RIPA: Radio-Immunoprecipitation Assay
BDM: 2,3-Butanedione monoxime
HBSS: Hanks’ Balanced Salt Solution
dPBS: Dulbecco’s Phosphate-Buffered Saline
Sqrt: Square root
FOV: field of view
FC: fold change
TBS: Tris-buffered saline
TBS-T: Tris-buffered saline, 0.1% Tween-20

## References

[1] C. Ménard, S. Pupier, D. Mornet, M. Kitzmann, J. Nargeot, P. Lory, Modulation of L-type calcium channel expression during retinoic acid-induced differentiation of H9C2 cardiac cells, Journal of Biological Chemistry 274 (1999) 29063–29070. 10.1074/jbc.274.41.29063.

[2] A.F. Branco, S.P. Pereira, S. Gonzalez, O. Gusev, A.A. Rizvanov, P.J. Oliveira, Gene expression profiling of H9c2 myoblast differentiation towards a cardiac-like phenotype, PLoS ONE 10 (2015) e0129303. 10.1371/journal.pone.0129303.

[3] C. Kankeu, K. Clarke, D. Van Haver, bcd Kris Gevaert, bc Francis Impens, bcd Anna Dittrich, H. Llewelyn Roderick, E. Passante, H.J. Huber, Quantitative proteomics and systems analysis of cultured H9C2 cardiomyoblasts during differentiation over time supports a “function follows form” model of differentiation, 14 (2018) 181. 10.1039/c8mo00036k.

[4] B.W. Kimes, B.L. Brandt, Properties of a clonal muscle cell line from rat heart, Experimental Cell Research 98 (1976) 367–381. 10.1016/0014-4827(76)90447-X.

[5] J. Hescheler, R. Meyer, S. Plant, D. Krautwurst, W. Rosenthal, G. Schultz, Morphological, biochemical, and electrophysiological characterization of a clonal cell (H9c2) line from rat heart, Circulation Research 69 (1991) 1476–1486. 10.1161/01.RES.69.6.1476.

[6] S.J. Watkins, G.M. Borthwick, H.M. Arthur, The H9C2 cell line and primary neonatal cardiomyocyte cells show similar hypertrophic responses in vitro, In Vitro Cell Dev Biol Anim 47 (2011) 125–131. 10.1007/s11626-010-9368-1.

[7] R.D. Harvey, J.W. Hell, CaV1.2 Signaling Complexes in the Heart, J Mol Cell Cardiol 58 (2013) 143–152. 10.1016/j.yjmcc.2012.12.006.

[8] C. Campero-Basaldua, J. Herrera-Gamboa, J. Bernal-Ramírez, S. Lopez-Moran, L.-A. Luévano-Martínez, H. Alves-Figueiredo, G. Guerrero, G. García-Rivas, V. Treviño, The retinoic acid response is a minor component of the cardiac phenotype in H9c2 myoblast differentiation, BMC Genomics 24 (2023) 431. 10.1186/s12864-023-09512-0.

[9] D. Avanzato, A. Merlino, S. Porrera, R. Wang, L.M. Munaron, D. Mancardi, Role of calcium channels in the protective effect of hydrogen sulfide in rat cardiomyoblasts, Cell Physiol Biochem 33 (2014) 1205–1214. 10.1159/000358690.

[10] X. Xu, L. Ruan, X. Tian, F. Pan, C. Yang, G. Liu, Calcium inhibitor inhibits high glucose-induced hypertrophy of H9C2 cells, Mol Med Rep 22 (2020) 1783–1792. 10.3892/mmr.2020.11275.

[11] A. Hirschy, F. Schatzmann, E. Ehler, J.-C. Perriard, Establishment of cardiac cytoarchitecture in the developing mouse heart, Developmental Biology 289 (2006) 430–441. 10.1016/j.ydbio.2005.10.046.

[12] P. Ahuja, P. Sdek, W.R. Maclellan, Cardiac Myocyte Cell Cycle Control in Development, Disease and Regeneration, Physiol Rev 87 (2007) 521–544. 10.1152/physrev.00032.2006.

[13] S.D. Lundy, W.-Z. Zhu, M. Regnier, M.A. Laflamme, Structural and Functional Maturation of Cardiomyocytes Derived from Human Pluripotent Stem Cells, Stem Cells Dev 22 (2013) 1991–2002. 10.1089/scd.2012.0490.

[14] M. Fritzsche, D. Li, H. Colin-York, V.T. Chang, E. Moeendarbary, J.H. Felce, E. Sezgin, G. Charras, E. Betzig, C. Eggeling, Self-organizing actin patterns shape membrane architecture but not cell mechanics, Nat Commun 8 (2017) 14347. 10.1038/ncomms14347.

[15] Y.M. Shirai, T.A. Tsunoyama, N. Hiramoto-Yamaki, K.M. Hirosawa, A.C.E. Shibata, K. Kondo, A. Tsurumune, F. Ishidate, A. Kusumi, T.K. Fujiwara, Cortical actin nodes: Their dynamics and recruitment of podosomal proteins as revealed by super-resolution and single-molecule microscopy, PLoS One 12 (2017) e0188778. 10.1371/journal.pone.0188778.

[16] F. Carton, S. Casarella, D. Di Francesco, E. Zanella, A. D’urso, L. Di Nunno, L. Fusaro, D. Cotella, M. Prat, A. Follenzi, F. Boccafoschi, Cardiac Differentiation Promotes Focal Adhesions Assembly through Vinculin Recruitment, Int J Mol Sci 24 (2023) 2444. 10.3390/ijms24032444.

[17] W.E. Louch, J.T. Koivumäki, P. Tavi, Calcium signalling in developing cardiomyocytes: implications for model systems and disease, J Physiol 593 (2015) 1047– 1063. 10.1113/jphysiol.2014.274712.

[18] S. Seki, M. Nagashima, Y. Yamada, M. Tsutsuura, T. Kobayashi, A. Namiki, N. Tohse, Fetal and postnatal development of Ca2+ transients and Ca2+ sparks in rat cardiomyocytes, Cardiovascular Research 58 (2003) 535–548. 10.1016/S0008-6363(03)00255-4.

[19] K.R. Gee, K.A. Brown, W.-N.U. Chen, J. Bishop-Stewart, D. Gray, I. Johnson, Chemical and physiological characterization of fluo-4 Ca2+-indicator dyes, Cell Calcium 27 (2000) 97–106. 10.1054/ceca.1999.0095.

[20] Z. Sun, T.C. Südhof, A simple Ca2+-imaging approach to neural network analyses in cultured neurons, J Neurosci Methods 349 (2021) 109041. 10.1016/j.jneumeth.2020.109041.

[21] M. Colpan, J. Iwanski, C.C. Gregorio, CAP2 is a regulator of actin pointed end dynamics and myofibrillogenesis in cardiac muscle, Commun Biol 4 (2021) 365. 10.1038/s42003-021-01893-w.

[22] J. Grego-Bessa, P. Gómez-Apiñaniz, B. Prados, M.J. Gómez, D. MacGrogan, J.L. de la Pompa, Nrg1 Regulates Cardiomyocyte Migration and Cell Cycle in Ventricular Development, Circ Res 133 (2023) 927–943. 10.1161/CIRCRESAHA.123.323321.

[23] W. Fu, Q. Liao, Y. Shi, W. Liu, H. Ren, C. Xu, C. Zeng, Transient induction of actin cytoskeletal remodeling associated with dedifferentiation, proliferation, and redifferentiation stimulates cardiac regeneration, Acta Pharm Sin B 14 (2024) 2537–2553. 10.1016/j.apsb.2024.01.021.

[24] W. Luo, Z.Z. Lieu, E. Manser, A.D. Bershadsky, M.P. Sheetz, Formin DAAM1 Organizes Actin Filaments in the Cytoplasmic Nodal Actin Network, PLoS One 11 (2016) e0163915. 10.1371/journal.pone.0163915.

[25] S. Xia, Y.B. Lim, Z. Zhang, Y. Wang, S. Zhang, C.T. Lim, E.K.F. Yim, P. Kanchanawong, Nanoscale Architecture of the Cortical Actin Cytoskeleton in Embryonic Stem Cells, Cell Reports 28 (2019) 1251–1267.e7. 10.1016/j.celrep.2019.06.089.

[26] M. Scarselli, P. Annibale, A. Radenovic, Cell Type-specific β2-Adrenergic Receptor Clusters Identified Using Photoactivated Localization Microscopy Are Not Lipid Raft Related, but Depend on Actin Cytoskeleton Integrity, J Biol Chem 287 (2012) 16768–16780. 10.1074/jbc.M111.329912.

[27] M.L. McCain, K.K. Parker, Mechanotransduction: the role of mechanical stress, myocyte shape, and cytoskeletal architecture on cardiac function, Pflugers Arch - Eur J Physiol 462 (2011) 89–104. 10.1007/s00424-011-0951-4.

[28] W.J. Kowalski, F. Yuan, T. Nakane, H. Masumoto, M. Dwenger, F. Ye, J.P. Tinney, B.B. Keller, Quantification of Cardiomyocyte Alignment from Three-Dimensional (3D) Confocal Microscopy of Engineered Tissue, Microscopy and Microanalysis 23 (2017) 826– 842. 10.1017/S1431927617000666.

[29] W.N. Arifin, W.M. Zahiruddin, Sample Size Calculation in Animal Studies Using Resource Equation Approach, Malays J Med Sci 24 (2017) 101–105. 10.21315/mjms2017.24.5.11.

[30] E. Ehler, T. Moore-Morris, S. Lange, Isolation and Culture of Neonatal Mouse Cardiomyocytes, J Vis Exp (2013) e50154. 10.3791/50154.

[31] D.A. Cantu, B. Wang, M.W. Gongwer, C.X. He, A. Goel, A. Suresh, N. Kourdougli, E.D. Arroyo, W. Zeiger, C. Portera-Cailliau, EZcalcium: Open-Source Toolbox for Analysis of Calcium Imaging Data, Front Neural Circuits 14 (2020) 25. 10.3389/fncir.2020.00025.

[32] P. Chomczynski, A reagent for the single-step simultaneous isolation of RNA, DNA and proteins from cell and tissue samples, Biotechniques 15 (1993) 532–534, 536–537.

[33] J. Vandesompele, K. De Preter, F. Pattyn, B. Poppe, N. Van Roy, A. De Paepe, F. Speleman, Accurate normalization of real-time quantitative RT-PCR data by geometric averaging of multiple internal control genes, Genome Biol 3 (2002) research0034.1-0034.11.

[34] O. Selmin, P.A. Thorne, P.T. Caldwell, P.D. Johnson, R.B. Runyan, Effects of trichloroethylene and its metabolite trichloroacetic acid on the expression of vimentin in the rat H9c2 cell line, Cell Biol Toxicol 21 (2005) 83–95. 10.1007/s10565-005-0124-3.

[35] J.E. Balmer, R. Blomhoff, Gene expression regulation by retinoic acid, Journal of Lipid Research 43 (2002) 1773–1808. 10.1194/jlr.R100015-JLR200.

[36] D. Wang, D. Zhao, Y. Li, T. Dai, F. Liu, C. Yan, TGM2 positively regulates myoblast differentiation via enhancing the mTOR signaling, Biochimica et Biophysica Acta (BBA) - Molecular Cell Research 1869 (2022) 119173. 10.1016/j.bbamcr.2021.119173.

[37] S. Schiaffino, A.C. Rossi, V. Smerdu, L.A. Leinwand, C. Reggiani, Developmental myosins: expression patterns and functional significance, Skeletal Muscle 5 (2015) 22. 10.1186/s13395-015-0046-6.

[38] M. Nemer, Genetic insights into normal and abnormal heart development, Cardiovascular Pathology 17 (2008) 48–54. 10.1016/j.carpath.2007.06.005.

[39] C. Cao, L. Li, Q. Zhang, H. Li, Z. Wang, A. Wang, J. Liu, Nkx2.5: a crucial regulator of cardiac development, regeneration and diseases, Front Cardiovasc Med 10 (2023) 1270951. 10.3389/fcvm.2023.1270951.

[40] S. Balasubramanian, S.K. Mani, H. Kasiganesan, C.C. Baicu, D. Kuppuswamy, Hypertrophic Stimulation Increases β-actin Dynamics in Adult Feline Cardiomyocytes, PLoS One 5 (2010) e11470. 10.1371/journal.pone.0011470.

[41] M. Suhaeri, R. Subbiah, S.Y. Van, P. Du, I.G. Kim, K. Lee, K. Park, Cardiomyoblast (H9c2) Differentiation on Tunable Extracellular Matrix Microenvironment, Tissue Engineering - Part A 21 (2015) 1940–1951. 10.1089/ten.tea.2014.0591.

[42] K.A. Taylor, D.W. Taylor, F. Schachat, Isoforms of α-Actinin from Cardiac, Smooth, and Skeletal Muscle Form Polar Arrays of Actin Filaments, J Cell Biol 149 (2000) 635–646.

[43] L.A. Swayne, N.P. Murphy, S. Asuri, L. Chen, X. Xu, S. McIntosh, C. Wang, P.J. Lancione, J.D. Roberts, C. Kerr, S. Sanatani, E. Sherwin, C.F. Kline, M. Zhang, P.J. Mohler, L.T. Arbour, A Novel Variant in the ANK2 Membrane-Binding Domain is Associated with Ankyrin-B Syndrome and Structural Heart Disease in a First Nations Population with a High Rate of Long QT Syndrome, Circ Cardiovasc Genet 10 (2017) e001537. 10.1161/CIRCGENETICS.116.001537.

[44] N.S. York, J.C. Sanchez-Arias, A.C.H. McAdam, J.E. Rivera, L.T. Arbour, L.A. Swayne, Mechanisms underlying the role of ankyrin-B in cardiac and neurological health and disease, Front Cardiovasc Med 9 (2022) 964675. 10.3389/fcvm.2022.964675.

[45] S.R. Cunha, P.J. Mohler, Ankyrin-based cellular pathways for cardiac ion channel and transporter targeting and regulation, Seminars in Cell and Developmental Biology 22 (2011) 166–170. 10.1016/j.semcdb.2010.09.013.

[46] C.F. Kline, J. Scott, J. Curran, T.J. Hund, P.J. Mohler, Ankyrin-B Regulates Cav2.1 and Cav2.2 Channel Expression and Targeting, J Biol Chem 289 (2014) 5285–5295. 10.1074/jbc.M113.523639.

[47] N. Chanthra, T. Abe, M. Miyamoto, K. Sekiguchi, C. Kwon, Y. Hanazono, H. Uosaki, A Novel Fluorescent Reporter System Identifies Laminin-511/521 as Potent Regulators of Cardiomyocyte Maturation, Sci Rep 10 (2020) 4249. 10.1038/s41598-020-61163-3.

[48] M. Tsikitis, Z. Galata, M. Mavroidis, S. Psarras, Y. Capetanaki, Intermediate filaments in cardiomyopathy, Biophys Rev 10 (2018) 1007–1031. 10.1007/s12551-018-0443-2.

[49] I. Shiraishi, D.G. Simpson, W. Carver, R. Price, T. Hirozane, L. Terracio, T.K. Borg, Vinculin is an Essential Component for Normal Myofibrillar Arrangement in Fetal Mouse Cardiac Myocytes, Journal of Molecular and Cellular Cardiology 29 (1997) 2041–2052. 10.1006/jmcc.1997.0438.

[50] A. Pott, S. Bock, I.M. Berger, K. Frese, T. Dahme, M. Keßler, S. Rinné, N. Decher, S. Just, W. Rottbauer, Mutation of the Na+/K+-ATPase Atp1a1a.1 causes QT interval prolongation and bradycardia in zebrafish, Journal of Molecular and Cellular Cardiology 120 (2018) 42–52. 10.1016/j.yjmcc.2018.05.005.

[51] T. Desplantez, Cardiac Cx43, Cx40 and Cx45 co-assembling: involvement of connexins epitopes in formation of hemichannels and Gap junction channels, BMC Cell Biology 18 (2017) 3. 10.1186/s12860-016-0118-4.

[52] B. Delorme, E. Dahl, T. Jarry-guichard, I. Marics, J.-P. Briand, K. Willecke, D. Gros, M. Théveniau-Ruissy, Developmental regulation of connexin 40 gene expression in mouse heart correlates with the differentiation of the conduction system, Developmental Dynamics 204 (1995) 358–371. 10.1002/aja.1002040403.

[53] C.W. Lo, A. Wessels, Cx43 Gap Junctions in Cardiac Development, Trends in Cardiovascular Medicine 8 (1998) 264–269. 10.1016/S1050-1738(98)00018-8.

[54] F. Jiang, K. Yin, K. Wu, M. Zhang, S. Wang, H. Cheng, Z. Zhou, B. Xiao, The mechanosensitive Piezo1 channel mediates heart mechano-chemo transduction, Nat Commun 12 (2021) 869. 10.1038/s41467-021-21178-4.

[55] D. Douguet, A. Patel, A. Xu, P.M. Vanhoutte, E. Honoré, Piezo Ion Channels in Cardiovascular Mechanobiology, Trends in Pharmacological Sciences 40 (2019) 956–970. 10.1016/j.tips.2019.10.002.

[56] T.M. da C.S. Carvalho, S. Cardarelli, M. Giorgi, A. Lenzi, A.M. Isidori, F. Naro, Phosphodiesterases Expression during Murine Cardiac Development, Int J Mol Sci 22 (2021) 2593. 10.3390/ijms22052593.

[57] H. Aoyama, Y. Ikeda, Y. Miyazaki, K. Yoshimura, S. Nishino, T. Yamamoto, M. Yano, M. Inui, H. Aoki, M. Matsuzaki, Isoform-specific roles of protein phosphatase 1 catalytic subunits in sarcoplasmic reticulum-mediated Ca2+ cycling, Cardiovascular Research 89 (2011) 79–88. 10.1093/cvr/cvq252.

