## Supplementary material for "Quantification of morphological, functional, and biochemical features of H9c2 rat cardiomyoblast retinoic acid differentiation": Figure S1

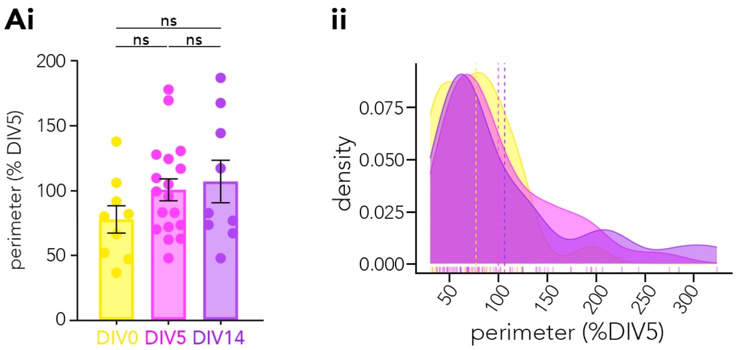


*Figure S1. H9c2 cell perimeter is not significantly altered during differentiation*. H9c2 cell morphology at DIV0, DIV5, and DIV14 were analyzed using the F-actin stain Phalloidin and nuclei were counted via Hoechst staining as in Figure 2. Perimeter was measured using a custom MATLAB script which resulted in no significant differences between groups (Ai) while the density distribution plots show a right shift for DIV5 and DIV14 compared to DIV0 (Aii). Results are graphed as Yellow = DIV0, Magenta = DIV5, Purple = DIV14. N = 3 replicates, n = 27 cells for all groups (3 cells from each of the 3 FOV were selected). ns, $p > 0.05$ by one-way ANOVA and post-hoc Tukey's multiple comparison and normalized to DIV5, see Table 1 for detailed statistics.
