## Supplementary material for "Quantification of morphological, functional, and biochemical features of H9c2 rat cardiomyoblast retinoic acid differentiation": Figure S2

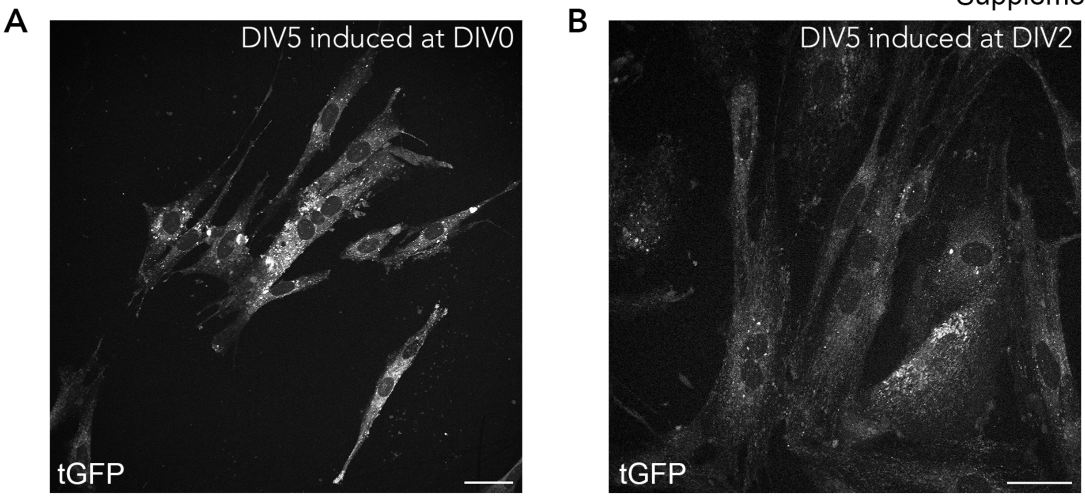


*Figure S1. H9c2 cell transfection has a high transfection efficiency.* H9c2 cells were seeded at 8.33 x 103 cells/cm2 and transfected with the Tet-inducible GFP plasmid the next day. Either the following day at (A) DIV0 or at (B) DIV2 GFP expression was induced with the addition of doxycycline hyclate, cells were differentiated until DIV5. Representative images show how efficient H9c2 transfection with jetPEI can be for future researchers investigating the role of genetic variants. DIV0, DIV2 induced (N =3, 4 FOV each).
