## Supplementary material for "Quantification of morphological, functional, and biochemical features of H9c2 rat cardiomyoblast retinoic acid differentiation": Table S1

Supplemental Table 1. Summary of genes analyzed across H9c2 cell differentiation

| gene | function | references |
| --- | --- | --- |
| *Rbp2* (retinol binding protein2) | facilitates dietary retinol uptake | increases with retinoic acid signaling [2,35] |
| *Tgm2* (transglutaminase2) | calcium dependent enzyme involved in myoblast differentiation | increases with retinoic acid signaling [8,35,36] |
| *Myh2* (myosin heavy chain2) | protein found in the thick sarcomeric filament | one of the major adult skeletal myosins [37] |
| *Tnnt2*, cTnT (cardiac troponin T) | anchors the troponin complex to tropomyosin. | highly expressed in cardiomyocytes, used in H9c2 literature as a marker of differentiated cells [2,3,8] |
| *Gata4* (GATA binding protein4) | zinc finger transcription factor involved in cardiomyocyte development | increases during cardiomyocyte development [38], levels remain unchanged during H9c2 cell differentiation 2025-04-25 12:53:00 PM |
| *Nkx2.5* (NK2 homeobox5) | transcription factor involved in cardiomyocyte development | one of the earliest markers of cardiac progenitor cells [39]. |
| *Actb*, β-actin | cytoskeletal protein important in cell migration | contributes to cardiomyocyte contractility [40] |
| *α-actinin* | cross links actin filaments | marker of cardiomyocytes [41,42] |
| ankyrin-B | scaffold protein linking ion channels to the cytoskeleton | scaffolds key cardiomyocyte proteins [43–46] |
| *Myom2* (Myomesin2) | key sarcomeric M band protein | increases with cardiomyocyte development [47] |
| *Vim*, vimentin | type III intermediate filament | not expressed in cardiomyocytes [48], decreases during H9c2 differentiation [34] |

| | | |
|---|---|---|
| vinculin | scaffold at cell-matrix adhesions | required for cardiomyocyte development [49] |
| *Atp1a1* (α-1 subunit of the $Na^+/K^+$ ATPase) | protein pump important in regulating ion balance in cardiomyocytes | important for ion regulation in repolarization [50] |
| *Cacna1c* (α-1C subunit of the voltage-dependent L-type $Ca^{2+}$ channel) | Calcium channel important for $Ca^{2+}$ import into cardiomyocytes | increases with H9c2 differentiation [1,8] |
| *Gja1*, Cx43 | gap junction protein | detected in the developing heart at E10.5 decreases after birth [51–53] |
| *Gja5* | major atrial gap junction protein | differentially regulated in heart development, increasing at P14 and then decreasing [51,52] |
| *Peizo1* | mechanosensitive ion channel | maintains normal heart function [54], expressed during embryonic heart development [55] |
| *Pde4a* (phosphodiesterase) | Key regulator of cAMP signalling in cardiomyocytes | Pde4a expression is stable across cardiomyocyte development [2,8,56] |
| *Ppp1ca* (α isoform of protein phosphatase1) | Negative regulator of $Ca^{2+}$ cycling | important for $Ca^{2+}$ regulation in adult cardiomyocytes [57] |
