## Supplementary material for "Quantification of morphological, functional, and biochemical features of H9c2 rat cardiomyoblast retinoic acid differentiation": Table S2

Supplemental Table 3. Western blot targets for characterizing cardiac protein expression.

| antibody | company, catalog | dilution | category |
|---|---|---|---|
| $\alpha$-actinin | Cell Signalling Technology, 3134S | 1:1000 | Cardiac-specific Markers |
| cTnT | Abcam, ab10214 | 1:2000 | |
| GATA4 | Santa Cruz Biotechnology, sc-25310 | 1:25 | Cardiac Transcription Factor |
| Nkx2.5 | Abcam, ab97355 | 1:400 | |
| $\beta$-actin | Sigma, A5441 | 1:2000 | Cytoskeleton |
| Vinculin | Sigma, MAB3574 | 1:60000, 1:10000 | |
| Vimentin | Cell Signalling Technology, 5741 | 1:80000 | |
| Connexin43 | Cell Signalling Technology, 3512 | 1:500 | Ion Channels |
| Peroxidase-Affinipure Donkey Anti-Mouse IGG (H+L) | Jackson Immuno Research, 715-035-150 | 1:8000 | Peroxidase Secondaries |
| Peroxidase-Affinipure Donkey Anti-Rabbit IGG (H+L) | Jackson Immuno Research, 711-035-152 | | |
| IRDye 800CW Donkey anti-Rabbit IgG (H + L) | Li-cor, 926-32213 | 1:8000 | Fluorescent Secondaries |
| IRDye 680RD Donkey anti-Mouse IgG (H + L) | Li-cor, 926-68072 | | |
| IRDye 800CW Donkey anti Mouse IgG (H + L) | Li-cor, 926-32212 | | |
| IRDye® 680RD Donkey anti-Rabbit IgG (H+L) | Li-cor, 926-68073 | | |
