## Supplementary material for "Quantification of morphological, functional, and biochemical features of H9c2 rat cardiomyoblast retinoic acid differentiation": Table S2

Supplemental Table 2. RT-qPCR assay panel for characterizing cardiac gene expression.

| gene | PrimeTime qPCR probe assay code | fluorophore | category |
|---|---|---|---|
| Rbp2 | Rn.PT.58.38085782 | Cy5 | Retinoic Acid Signaling |
| Tgm2 | Rn.PT.58.13093079 | Cy5 | |
| Myh2 | Mm.PT.58.7469087 | Cy5 | Cardiac-specific Markers |
| Tnnt2 | Mm.PT.58.12681437 | Hex | |
| Ctnnb1 | Mm.PT.58.10078649 | Cy5 | Cardiac Transcription Factor |
| GATA4 | Mm.PT.58.30024990 | Fam | |
| Nkx2.5 | Rn.PT.58.11528139 | Fam | |
| Actb | Mm.PT.39a.22214843.g | Fam | Cytoskeleton |
| Actn1 | Mm.PT.58.30787926 | Hex | |
| Myom2 | Rn.PT.58.10611775 | Hex | |
| Vim | Mm.PT.58.8720419 | Fam | |
| Atp1a1 | Rn.PT.58.17975975 | Fam | Ion Channels |
| Cacna1c | Rn.PT.58.16361922 | Hex | |
| GJA1 | Rn.PT.58.10553710 | Cy5 | |
| GJA5 | Rn.PT.58.7544937 | Fam | |
| Piezo1 | Rn.PT.58.46120084 | Cy5 | |
| Polr2a | Mm.PT.39a.22214849 | Cy5 | Housekeeping |
| Ppia | Mm.PT.39a.2.gs | Fam | |

| | | | |
|---|---|---|---|
| Pde4a | Rn.PT.58.6046707 | Cy5 | Housekeeping |
| PPP1CA | Mm.PT.58.23629119 | Hex | |
